# Context-Dependent Probability Estimation and its Neurocomputational Substrates

**DOI:** 10.1101/624163

**Authors:** Wei-Hsiang Lin, Justin L. Gardner, Shih-Wei Wu

## Abstract

Many decisions rely on how we evaluate potential outcomes associated with the options under consideration and estimate their corresponding probabilities of occurrence. Outcome valuation is subjective as it requires consulting internal preferences and is sensitive to context. In contrast, probability estimation requires extracting statistics from the environment and therefore imposes unique challenges to the decision maker. Here we show that probability estimation, like outcome valuation, is subject to context effects that bias probability estimates away from other stimuli present in the same context. However, unlike valuation, these context effects appeared to be scaled by estimated uncertainty, which is largest at intermediate probabilities. BOLD imaging showed that patterns of multivoxel activity in dorsal anterior cingulate cortex (dACC) and ventromedial prefrontal cortex (VMPFC) predicted individual differences in context effects on probability estimate. These results establish VMPFC as the neurocomputational substrate shared between valuation and probability estimation and highlight the additional involvement of dACC that can be uniquely attributed to probability estimation. As probability estimation is a required component of computational accounts from sensory inference to higher cognition, the context effects found here may affect a wide array of cognitive computations.

**Highlights:**

1. Context impacts subjective estimates on reward probability – Stimuli carrying greater variance are more strongly affected by other stimuli present in the same context
2. This phenomenon can be explained by reference-dependent computations that are gated by reward variance
3. Multivoxel patterns of dACC and VMPFC activity predicts individual differences in context effect on probability estimate

## INTRODUCTION

When we evaluate a potential reward, such as a job offer, other offers on the table form a unique context that shapes our expectations and influences the way we feel about it. Psychologists and economists model such expectation as a reference point and potential rewards (e.g. different job offers) as gains or losses relative to the reference point (Kahneman & Tversky, 1979; Kőszegi & Rabin, 2006). Such context-dependent valuation impacts many decisions we make and is not only observed in humans, but also in many other species such as rats (Belke, 1992; Gallistel, 2000), birds (Pompilio & Kacelnik, 2010) and non-human primates (Tremblay & Schultz, 1999; Chen et al., 2006; Zimmerman et al., 2018).

However, to make decisions in uncertain environments, organisms not only face the task of evaluating potential rewards but also need to estimate their probabilities of occurrence (Bernoulli, 1738/1954; von Neumann & Morgenstern, 1944). Understanding how people use probability information has received considerable attention and people are known to subjectively weight probability information rather than using objective probability information when making decisions: they tend to overweight small probabilities but underweight moderate to large probabilities. (Kahneman & Tversky, 1979; Tversky & Kahneman, 1992; Wu & Gonzalez, 1996; Fox & Tversky, 1998; Gonzalez & Wu, 1999; Hertwig & Erev, 2009; Wu et al., 2009).

But is probability estimation — like valuation — context-dependent? Going back to the job-offer example, when one is instead estimating the probability of receiving a potential offer, if there is another offer that is relatively very likely or very unlikely to happen, would it affect how she or he estimates the probability of getting this particular offer? Probability estimate of an event may be affected by other events that take place close in time, which makes it susceptible to context effect in a way similar to how context impacts valuation. However, probability estimation may be different from valuation in that very unlikely or very likely events are less variable than those carrying intermediate probabilities, so that context effect on probability estimation may not be uniform in magnitude across probabilities. Despite such intuitive appeal, standard models of decision making under uncertainty and risk typically do not consider probability estimation to be context-dependent (Bernoulli, 1738/1954; von Neumann & Morgenstern, 1944; Kahneman & Tversky, 1979; Fox & Tversky, 1998) and context effects on probability estimation are seldom investigated.

At the neural implementation level, we consider whether probability estimation would be computed by shared or disassociated cortical structures for value estimation. The first hypothesis of shared cortical implementation centers on the similarity in computations between valuation and probability estimation. That is, if there are context effects on probability estimation, then the similarity with respect to valuation might suggest that both are implemented by the same neural system. In this case, the orbitofrontal cortex (OFC) and ventromedial prefrontal cortex (VMPFC) are candidate regions for context-dependent probability estimation, as accumulating evidence indicate their involvement in subjective-value computations (Kable & Glimcher, 2009; Bartra et al., 2013; Clithero & Rangel, 2014), in particular the relative and context-dependent representations of subjective value found in these regions (Tremblay & Schultz, 1999; Elliot et al., 2008; Padoa-Schioppa, 2009; Palminteri et al., 2015; Yamada et al., 2018). By contrast, the second hypothesis of disassociated cortical structures starts from the potential context effects in probability estimation that are unique to valuation and thus points to neural systems dissociable from valuation. In this case, regions specialized for coding uncertainty and extracting summary statistics, such as mean and variance, from reward history should be highly involved. Previous studies indicate that the dorsal anterior cingulate cortex (dACC) would be the candidate region as it had been shown to represent uncertainty-related statistics (Behrens et al., 2007; Christopoulos et al., 2009) and engage in tracking and comparing between recent and distant reward history (Wittman et al., 2016; Kolling et al., 2016).

To investigate how context impacts probability estimation at the behavioral, computational and neural implementation levels, in a simple stimulus-reward association task, human subjects were asked to estimate probability of reward associated with visual stimuli through experience. Context was manipulated by pairing stimuli carrying different probabilities of reward in different blocks of trials. We found that, similar to valuation, context effect on probability estimation was reference-dependent. However, unlike valuation, it was scaled by the uncertainty of reward outcomes such that when there is larger uncertainty on which potential outcome would occur (e.g. 50/50 of reward or no reward) there was greater context effect than stimuli with smaller standard deviation (10% or 90% reward). Unexpectedly, imaging results showed that both valuation and uncertainty-coding regions are involved in context-dependent probability estimation. Together, these results point to common neurocomputational substrates shared between probability estimation and valuation but highlight the additional involvement of uncertainty-coding regions that are unique in probability estimation — a central and important cognitive function relevant to a wide array of contemporary problems within neuroscience (Knill & Richards, 1996; Tenenbaum et al., 2011; Pouget et al., 2013).

## RESULTS

In the MRI scanner, subjects (n=34) performed a stimulus-reward association task where they were asked to estimate reward probability, but no choice was required (Fig. 1A). On each trial, subjects were presented with an abstract visual stimulus that carried a unique probability of reward and asked to estimate its reward probability. After probability estimation, feedback of whether subjects won a monetary reward was provided. In a block of trials, subjects would repeatedly face two different stimuli that appeared in random order. Context was defined by the two stimuli the subjects encountered in a block of trials and was manipulated such that stimuli carrying the same probability of reward were experienced in two different contexts that differed in the probability of reward associated with the other stimulus present in the context (Fig. 1B). For example, for 50% reward (middle column in Fig. 1B), one stimulus was paired with a stimulus carrying 10% reward (context 1, first row in Fig. 1B) and the other with a 90% reward (context 3, third row in Fig. 1B). With this design, we were able to manipulate context independently of probability of reward (10%, 50%, and 90%). In Fig. 1C we illustrate the three different contexts used in this experiment with example trial ordering. The goal of the experiment was to investigate how context impacts probability estimate by comparing probability estimate of the stimuli carrying the same reward probability but experienced in different contexts. For example, we want to compare 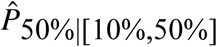 (probability estimate of the stimulus carrying 50% reward in context 1 where the other stimulus had 10% reward) with 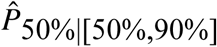 (probability estimate of the stimulus carrying 50% reward in context 3 where the other stimulus had 90% reward).

**Figure 1.**
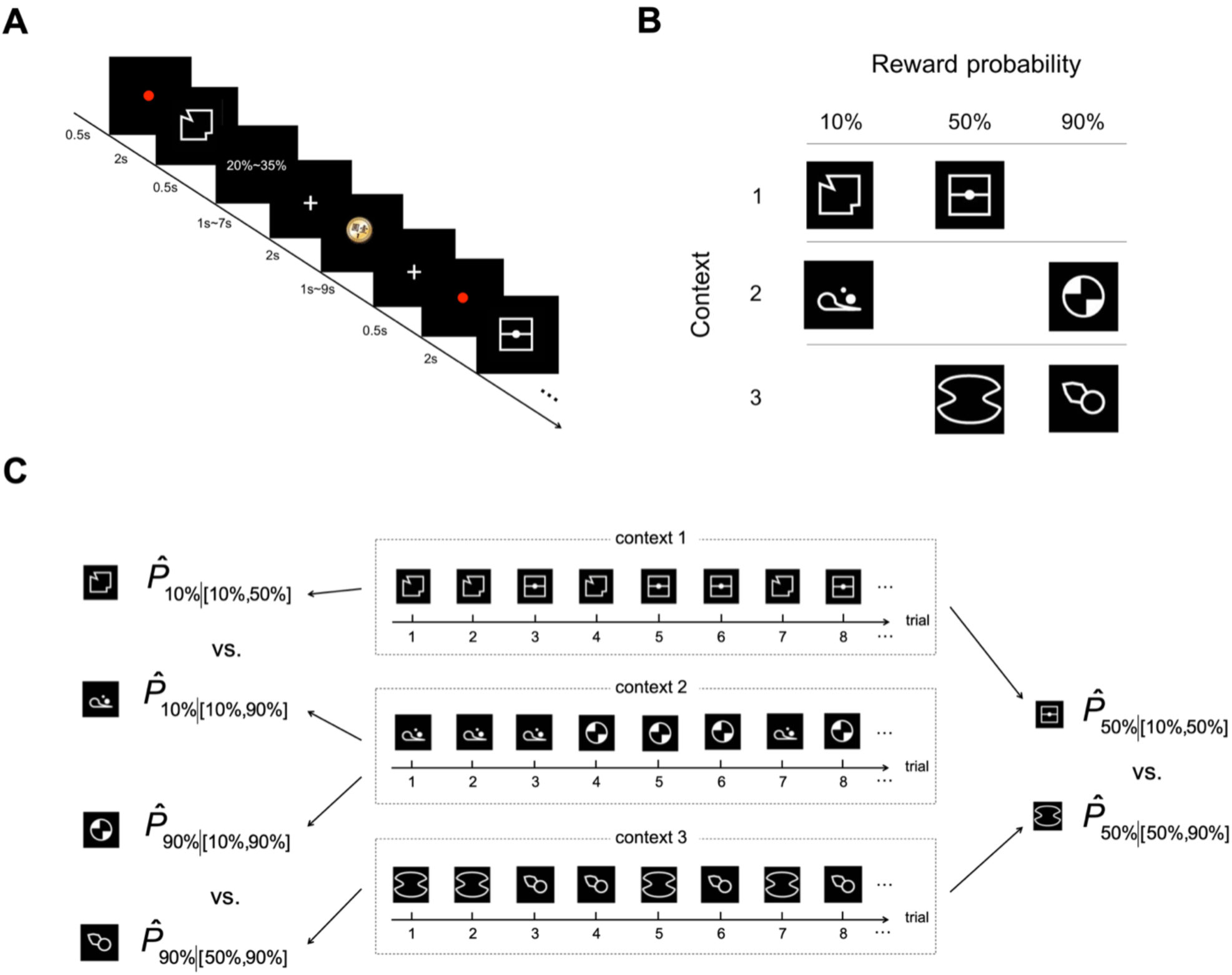
Task design. **A.** Trial sequence. On each trial, subjects were presented with an abstract visual stimulus and had to indicate his/her estimate of the probability of reward associated with it with a button-press. A brief feedback (0.5 sec) on the probability estimate subjects gave was then provided. After a variable inter-stimulus interval (ISI, 1∼7 sec), subjects received feedback on whether she/he received a monetary reward. A variable inter-trial interval (ITI, 1∼9 sec) was implemented before the start of a new trial. In a block of trials, two stimuli each carrying a unique probability of reward appeared repeatedly in random order. **B.** Manipulation of context. Context was defined by the stimuli that appeared in a block of trials. There were three contexts (rows in the table), each consisting of two visual stimuli carrying different probabilities of reward. For each probability, there were two different stimuli assigned to it. The two stimuli with the same probability were experienced under two different contexts. This design allowed us to compare how probability estimate was affected by context. **C.** Example trial ordering of the three contexts. The goal of the experiment was to investigate context effect on probability estimate, i.e. compare probability estimate of stimuli carrying the same reward probability but experienced in different contexts.

### Context effects on probability estimate

We found significant context effect on probability estimate when reward probability was 50%, but not at 10% (t=1.9419, df=33, p=0.0607) or 90% (t=1.9441, df=33, p=0.0605). Subjects gave larger estimates when the 50%-reward stimulus was experienced with a 10%-reward stimulus than with a 90%-reward stimulus (t=5.3195, df=33, p<0.001), as indicated by the two bars in the center of Fig. 2A — the bar on the left indicates probability estimate of the 50%-reward stimulus (S) when the other stimulus (OS) in the context was 10%; the bar on the right indicates 50% probability estimate when OS in the context was 90%. Looking at estimates of individual subjects, 27 out of 34 subjects’ estimates were larger in the [10%,50%] context than the [50%,90%] context. This can be captured by the direction of tilt of the black lines (each representing a single subject’s mean probability estimates) (Fig. 2A), showing that most subjects gave higher estimates to the 50% stimulus when it was paired with a 10%-reward stimulus than with a 90%-reward stimulus. Importantly, we observed this effect despite the fact that frequency of reward the subjects experienced was the same between different contexts (two bars in the center of Fig. 2B). This indicates that, for the 50%-reward stimuli, context effect on probability estimate was not driven by its own reward history.

**Figure 2.**
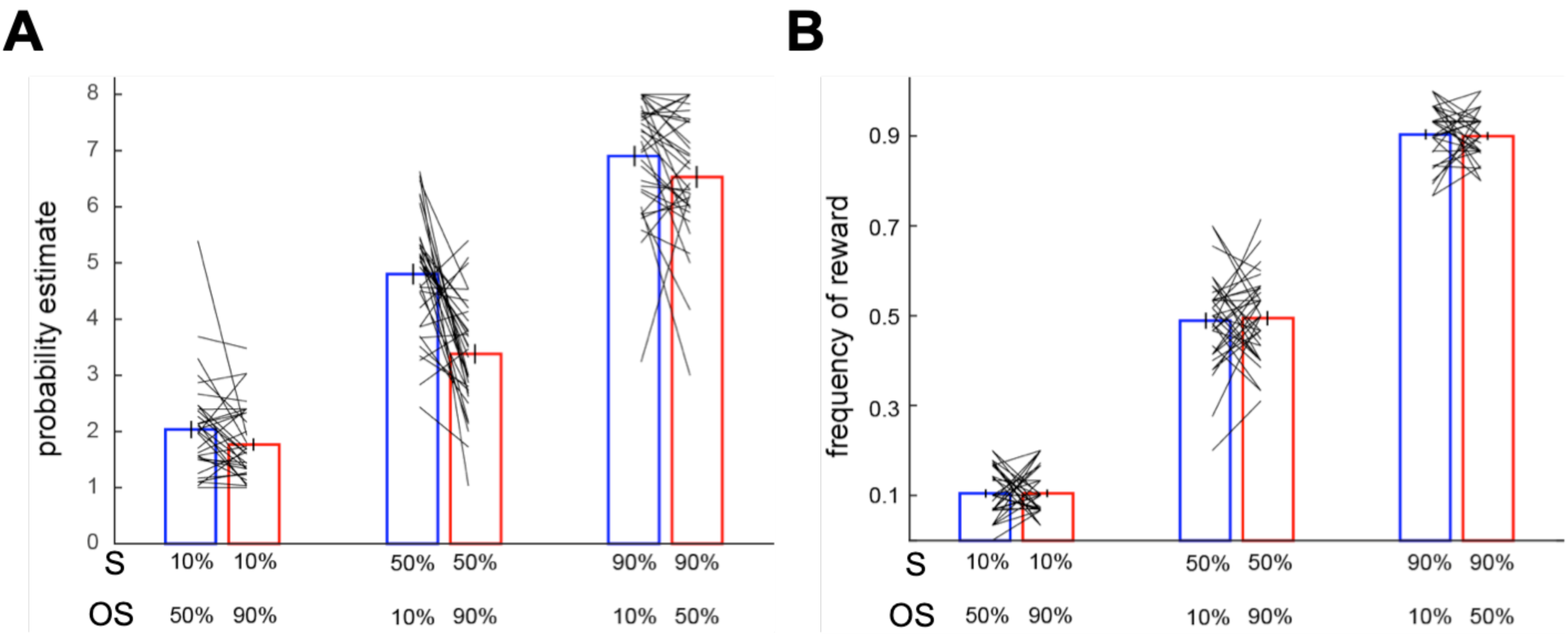
Context effect on probability estimate. **A.** Subjects’ estimates on probability of reward associated different visual stimuli. The stimuli carrying the same reward probability are plotted together and compared using paired t-test. The bars represent the mean probability estimates (across subjects) and the tilted black lines represent individual subjects’ data. **B.** Frequency of reward the subjects experienced during the experiment. Conventions are the same as in **A.** Error bars represent ±1 SEM.

It is important to note that, even though context effect was not statistically significant at 10% and 90%, we did observe a trend: the probability estimate of a stimulus was inversely related to the probability of reward associated with the other stimulus present in the context. At 10% reward, subjects tended to give smaller estimate when the other stimulus carried a 90% than 50% reward. At 90% reward, subjects tended to give larger estimate when the other stimulus carried a 10% than 50% reward.

### Context-dependent probability estimate predicts choice behavior

We also found that context effect on probability estimate further predicted subjects’ choice behavior. In a lottery decision task (a behavioral session) that followed the probability estimation task (fMRI session), subjects on each trial faced two stimuli they encountered in the fMRI session and were asked to choose the one they preferred so that one of their choices would be selected at random and realized to determine his/her payoff. Each stimulus they faced carried the same reward probability as in the fMRI session and the reward magnitude was fixed across all stimuli so that subjects should pick the one she or he believed to have the larger probability of reward. Each pair was presented multiple times so that we could calculate choice probability (the fraction of trials in which subjects chose one lottery over the other) for each subject. We analyzed choice probability on the three pairs of stimuli each consisting of stimuli carrying the same probability of reward but were experienced under different contexts. Subjects’ choice behavior was predicted by the context-dependent probability estimate: they preferred the 50%-reward stimulus with larger probability estimate (in the [10%,50%] context) to the one with smaller estimate (in the [50%,90%] context) (middle bar in Fig. 3) (t(32)=3.9389, p<0.0005). In contrast, for the 10% and 90% pairs, subjects were indifferent between the stimuli carrying the same probability of reward (left and right bars in Fig. 3; 10%-reward pair: t(32)=1.473, p=0.15; 90%-reward pair: t(32)=1.7426, p=0.09) just as their probability estimates did not differ between contexts. However, it is worth noting that the direction of choice in those pairs was still qualitatively consistent with the probability estimates (Fig. 2A): subjects tended to chose more often the stimuli that with larger probability estimate. In summary, these results demonstrate that the effect of context effect was not only confined to subjects’ probability estimate – it further impacted subjects’ preference for the stimuli.

**Figure 3.**
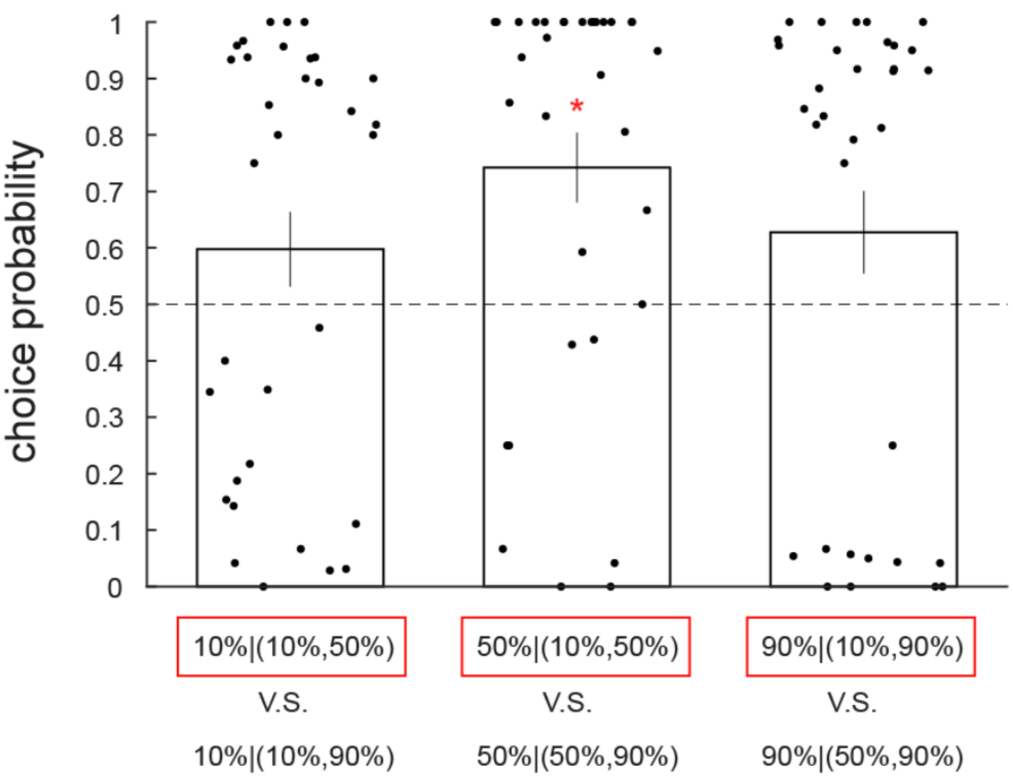
Choice probability in the post-fMRI lottery decision task. Three pairs of options, each representing a choice between stimuli carrying the same reward probability but experienced in different contexts, are shown. For each pair, we plot the choice probability of the option highlighted in red. The bar indicates the mean choice probability averaged across all subjects. Error bars represent ±1 SEM. Each data point represents choice probability of a single subject. For each pair, we tested whether the mean choice probability is different from 50%. The red star symbol indicates significant difference at p<0.05.

### Dynamics of probability estimate and response time

We examined trial-by-trial probability estimate and found that context effect on 50% reward emerged relatively early and persisted throughout the experiment (Fig. 4B). In Fig. 4, we plot the trial-by-trial mean probability estimate and the mean response time (both averaged across subjects) under different contexts. Another aspect of the dynamics worth mentioning is that subjects’ probability estimate clearly deviated from the actual frequency of reward that they experienced (the red and blue step-function traces in Fig. 4B). Compared with frequency of reward, subjects underestimated the 50% probability of reward (blue curve) when it was experienced with a 90%-reward stimulus in the [50%, 90%] context but overestimated 50% when it was experienced with a 10%-reward stimulus in the [10%,50%] context. By contrast, we did not observe these patterns on the 10%-reward (Fig. 4A) and 90%-reward stimuli (Fig. 4C). Qualitatively, however, we did see that subjects gave larger estimates to the 10%-reward stimulus when it was experienced with a 50%-reward stimulus (red in Fig. 4A) than the 10% stimulus experienced with a 90%-reward stimulus (blue in Fig. 4A). For 90% reward, even though not statistically significant, subjects gave larger estimates to the 90%-reward stimulus in the [10%,90%] context (red in Fig. 4C) than in the [50%,90%] context (blue in Fig. 4C).

**Figure 4.**
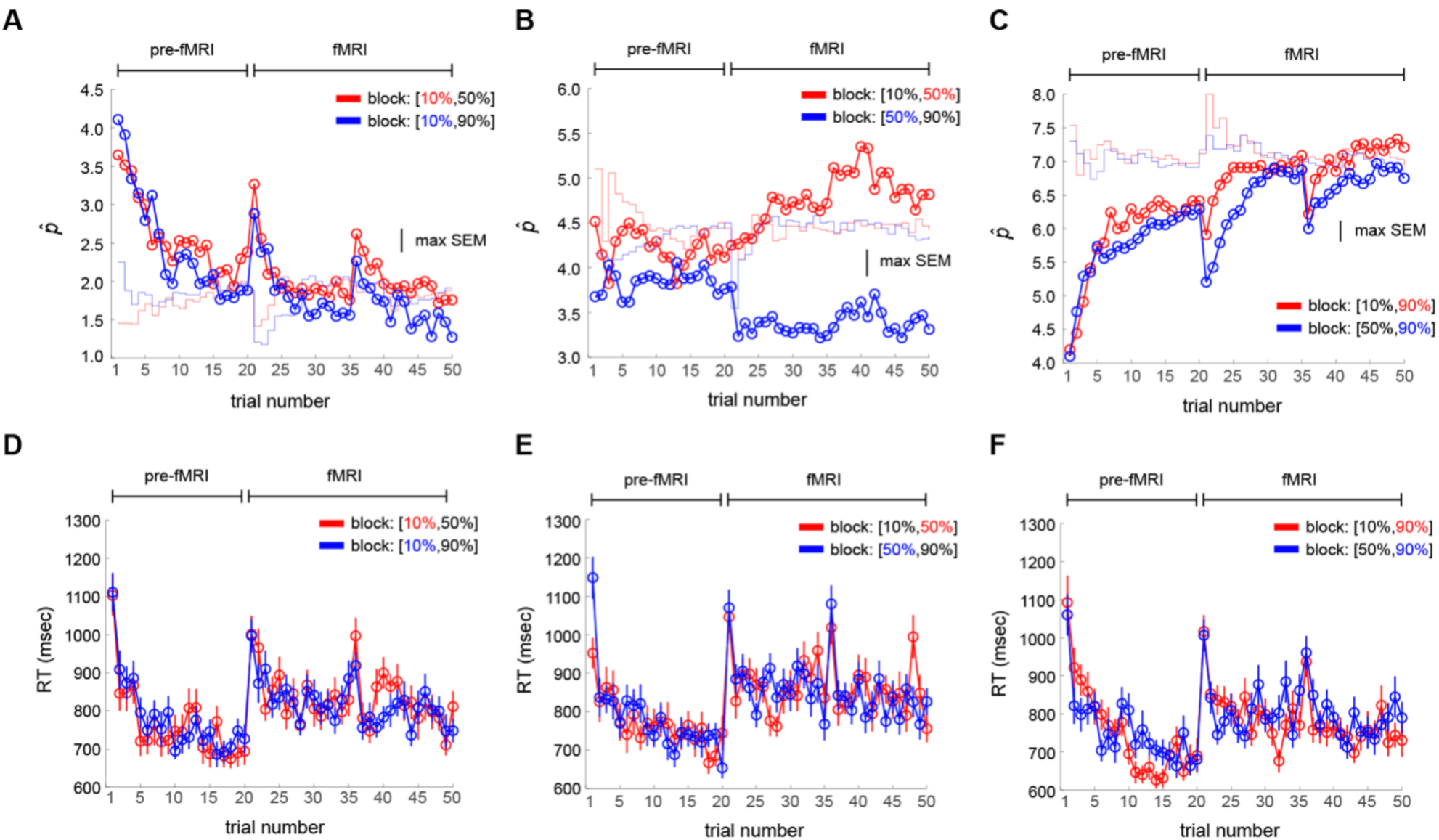
Comparison of trial-by-trial probability estimate and response time (RT) between contexts. Data included the pre-fMRI practice session and the fMRI session. **A-C.** Probability estimate. The scale on the y-axis indicates interval estimate (from 1 to 8 with 1 representing the smallest probability interval [0%,5%] and 8 representing the largest probability interval [95% 100%]). **A.** 10%-reward stimuli. Each data point represents the mean probability estimate of a single trial. Red: 10% stimulus in the [10%,50%] context. Blue: 10% stimulus is in the [10%,90%] context. The colored step function trace represents the frequency of reward experienced by the subjects. **B.** Comparison of the 50%-reward stimuli. Red: 50% stimulus in the [10%,50%] context. Blue: 50% stimulus in the [50%,90%] context. **C.** Comparison of the 90%-reward stimuli. Red: 90% stimulus in the [10%,90%] context. Blue: 90% stimulus in the [50%,90%] temporal context. **D-F.** RT for 10% stimuli (D), 50% stimuli (E) and 90% stimuli (F). Conventions are the same as in **A-C**. Error bars represent ±1 SEM. The bumps on trial number 21 and 36, seen on A, C, D-F, reflect the beginning of a new block of trials.

In contrast to the probability estimate, we did not find context effect on response time (RT) – the time it took the subjects to indicate probability estimate – in all three probabilities of reward (Fig. 4D-F). Together with the data on probability estimates, these results indicate that context effect on 50%-reward probability estimate was stable throughout trials in the fMRI session and that context effect on probability estimate was not reflected on response time data.

### A behavioral control experiment replicated context effect on probability estimate

We conducted an additional behavioral control experiment (n=20 subjects) to address a potential confound on the assignment of buttons to indicate probability estimate. In the experiment, because the boundary between the two buttons for small probabilities was set at 5% and at 95% for large probabilities instead of 10% and 90% respectively (see *Methods* for details), it is possible that non-significant context effect at 10% and 90% was due to lack of sensitivity on the interval estimate to detect differences in probability estimates at 10% and 90% between different contexts. In the control experiment, a total of 10 buttons were used, each representing a 10% interval and therefore addressed the above concern (see *Behavioral control experiment* in *Methods*). We replicated the results observed in the main experiment: For the 50%-reward stimuli, subjects gave larger estimates when 50% was paired with 10% than with 90% (p<0.05). For the 10% and 90% stimuli, no significant context effect was found (Fig. 5A-C). Furthermore, as in the main experiment, context effects on probability estimate predicted subjects’ choice behavior in the lottery decision task that followed the probability estimation task (Fig. 5D). Subjects chose more often the stimulus assigned with larger probability estimate when its reward probability was 50% (t=2.9876, df=19, p<0.01). By contrast, when stimulus reward probability was 10% or 90%, there was no significant difference in choice probability (10% reward: t=1.2652, df=19 p=0.2211; 90%: t(19)=0.7771, p=0.4467).

**Figure 5.**
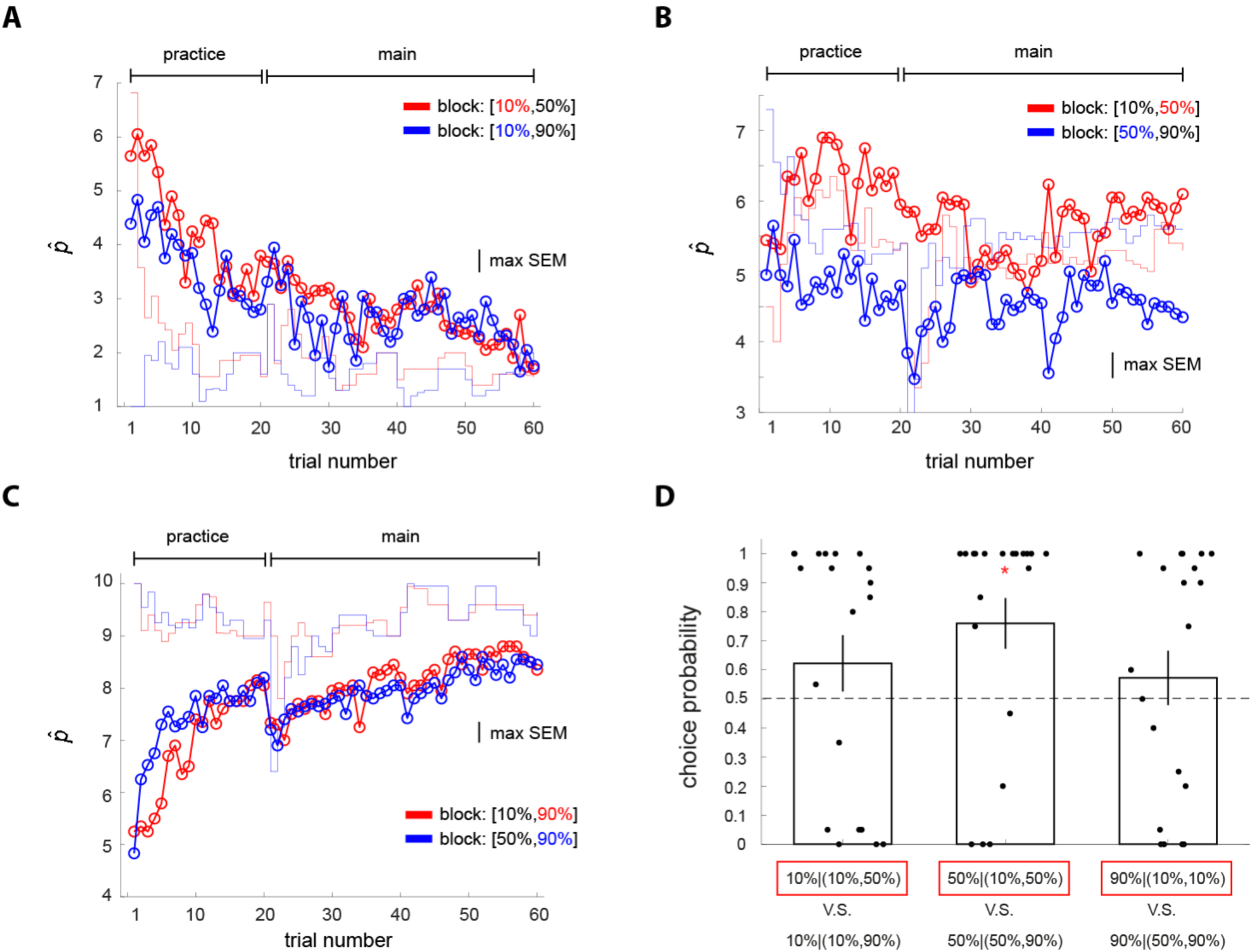
Behavioral control experiment. Conventions are the same as in Figure 3 and 4. **A-C.** Probability estimate. The scale on the y-axis indicates interval estimate (from 1 to 10 with 1 representing the smallest probability interval [0%,10%] and 10 representing the largest probability interval [90% 100%]).

### Context effect on probability estimate is driven by reference and variance dependencies

To systematically explain the context effects, we developed and tested a mathematical model for context-dependent probability estimation (see *Methods* for details; Fig. 6A for an illustration). In short, the model proposes that when computing probability estimate of a stimulus 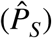, the decision maker uses the frequency of reward (*f_S_*) she or he experienced but is biased by the overall frequency of reward (*f*_overall_) associated with a context. Here (*f*_overall_) is the frequency of reward averaged across different stimuli in a context. The bias arises from treating (*f*_overall_) as a reference point and comparing *f_S_* with it (*f_S_* − *f*_overall_). For example, for a stimulus carrying 50% reward, its *f_S_* would be greater than *f*_overall_ in the [10%,50%] context where the other stimulus carries a 10% reward but would be smaller than *f*_overall_ in the [50%,90%] context where the other stimulus carries a 90% reward. The model therefore predicts that subjects would give larger probability estimate to the stimulus carrying 50% reward in the [10%,50%] context than in the [50%,90%] context. However, simply having the bias term (*f_S_* − *f*_overall_) alone cannot explain why we observed context effect at 50% but did not find significant context effect at 10% and 90% reward. We reasoned that this is due to the level of uncertainty regarding which outcome (reward or no reward) would occur: when the level of uncertainty is higher (e.g. at 50% reward), the impact of (*f_S_* − *f*_overall_) on probability estimate is larger than when the level of uncertainty is lower (e.g. at 10% or 90% reward). This is modeled by having (*f_S_* − *f*_overall_) weighted by the level of uncertainty, which is modeled by either the variance 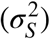 or standard deviation (*σ_S_*) of potential outcomes associated with the stimulus. Below is a version of the model that uses (*σ_S_*) to weight the impact of (*f_S_* − *f*_overall_).

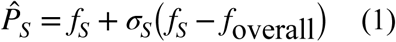

**Figure 6.**
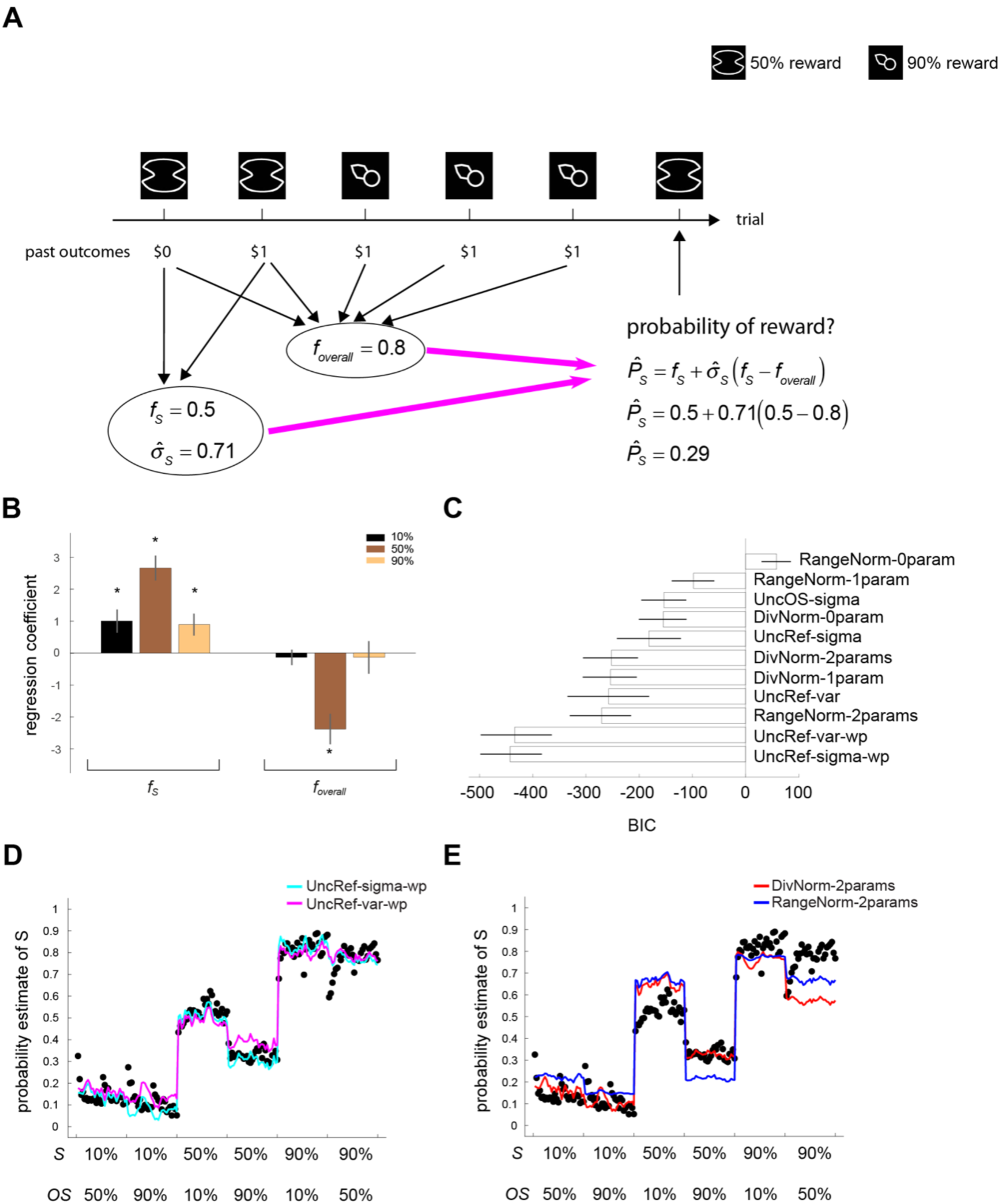
Computational models for context-dependent probability estimation. **A.** The context-dependent model for probability estimate developed in this study. **B.** Regression analysis. In three separate multiple regression models each examining a particular reward probability, we estimated the weight subjects assigned to frequency of reward associated with a stimulus (*f_S_*) and the overall frequency of reward associated with the context (*f*_overall_) and plot the mean regression coefficient (across all subjects). Error bars represent ±1 SEM. The star symbol indicates statistical significance (testing the mean regression coefficient against 0 at p<0.05). **C.** Model comparison. We fitted 11 different models and plot the value of their respective Bayesian information criterion (BIC). Smaller BIC values indicate better models. The best models are two versions of the context-dependent model developed in this study (UncRef-var-wp, UncRef-var-wp). The range of error bar represents 95% confidence interval. **D.** Model fitting results of best two models (UncRef-sigma-wp and UncRef-var-wp). **E.** Model fitting results of divisive normalization model (DivNorm-2params) and range normalization model (RangeNorm-2params). Each data point (black) in **D** and **E** represents the mean probability estimate (across all subjects) of a particular trial order. *S*: stimulus of interest. *OS*: the other stimulus in the same context.

Equation (1) can be rewritten as 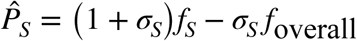, which made several predictions on the weights assigned to *f_S_* and *f*_overall_ by the subjects that were confirmed by multiple regression analysis on subjects’ probability estimates using *f_S_* and *f*_overall_ as regressors (Fig. 6A). First, the regression analysis confirmed that the regression coefficient of *f_S_* was positive and significantly different from 0 for all three different reward probabilities. Second, the regression analysis confirmed that coefficient of *f_S_* associated with 50% reward was greater than that associated with 10% and 90% reward. Third, the regression analysis confirmed that coefficient of *f*_overall_ associated with 50% reward was negative and its magnitude was larger than that associated with the 10%-reward and 90%-reward stimuli. Together, these results provide support to our model for context-dependent probability estimation.

### Model comparison

We further compared our model with normalization models that have been useful in describing context-dependent valuations (Louie et al., 2011, 2013; Yamada et al., 2018). Here we consider two classes of normalization models, the divisive normalization model (DivNorm) and the range normalization model (RangeNorm). Divisive normalization computes probability estimate of a stimulus by having its frequency of reward divided by the sum of the frequency of reward associated with the two stimuli in a context (Eq. 8 in *Methods*), while range normalization uses the difference between the maximum and the minimum stimulus reward frequency in a context in the denominator to compute probability estimate (Eq. 9 in *Methods*). Both model classes predict large context effect on probability estimate at 50% reward and non-significant effect at 10%, which are consistent with what we observed in this dataset. However, both models also predict large context effect at 90% reward, which we did not observe in subjects’ probability estimate. It is also worth mentioning that, in contrast to our model, these models do not use information about reward variance or standard deviation to compute probability estimates.

A total of 11 different models – five versions of our model, three versions of DivNorm, and three versions of RangeNorm – were fitted to the trial-by-trial mean probability estimates (across subjects) using method of maximum likelihood (see *Methods*). To compare these models, we computed Bayesian information criterion (BIC) and for each model estimated its 95% confidence interval using non-parametric Bootstrap method (Efron & Tibshirani, 1994). The best models are two versions of our model (Fig. 6B) – the uncertainty-weighted reference-dependent model that uses standard deviation as a measure of uncertainty and that considers overestimation of small probability and underestimation of large probability (UncRef-sigma-wp) and the same model except using variance instead as a measure of uncertainty (UncRef-var-wp). In Fig. 6C, we show the model fits of these two models. The maximum likelihood estimate of the parameters were applied to the models to generate the model-predicting curves (color-coded), each describing a model. To further compare these model fits with normalization models, in Fig. 6D, we show the model fits of two normalization models, one from divisive normalization (DivNorm-2params) and the other from range normalization (RangeNorm-2params). Both models fail to capture non-significant context effect at 90% reward. For 50% reward, even though both models can describe the directions of context effect, the fits were visibly worse compared with our models (Fig. 6C).

In summary, the best models have three features. First, it is comparative in the sense that the probability estimate of a stimulus comes from comparing its frequency of reward with a reference point that is the average frequency of reward associated with the context the stimulus is in. Second, this comparison effect (or reference-dependent effect) is weighted by the uncertainty on which outcome would occur associated with the stimulus such that the effect is stronger for stimuli carrying larger uncertainty (having larger variance/standard deviation). Finally, the best models capture overestimation of small probabilities and underestimation of large probabilities, which were also observed in Fox and Tversky (1998).

### fMRI results

We performed both univariate general linear model (GLM) analysis and multivoxel pattern analysis (MVPA) on the fMRI data. All the analyses focused on two time windows during a trial – when the visual stimulus was presented and subjects had to estimate probability (stimulus presentation) and when the reward outcome (whether subjects received a reward or not reward) was shown (reward feedback).

### Average brain activity did not represent context effect on probability estimate

We found no evidence for context effect based on average brain activity. First, at stimulus presentation, for each reward probability separately, we compared activity between different contexts and got null results (p>0.05 whole-brain FWE corrected for multiple comparisons using Gaussian random field theory with cluster-defining threshold at z>2.3). Second, we examined whether there were regions representing individual differences in context effect on 50%-reward (the probability that showed significant context effect at the behavioral level). For each subject separately, we computed the difference in the mean probability estimate between contexts 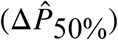 and used it as a behavioral measure of context effect. We performed a group-level analysis using 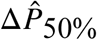 as a covariate and found no brain region whose activity difference between contexts correlated with individual differences in context effect (p>0.05 whole-brain FWE corrected for multiple comparisons using Gaussian random field theory with cluster-defining threshold at z>2.3). Finally, region-of-interest (ROI) analysis on the valuation network in the ventromedial prefrontal cortex (VMPFC) and ventral striatum (VS) (using coordinates from Clithero & Rangel, 2014) also did not show significant result on the two analyses described above. Together, these findings indicate that context effect on probability estimate is not represented by brain activity averaged across trials at the time of stimulus presentation. For results at the time of reward feedback, see *Replication of reward prediction error signal and reward magnitude in VMPFC and striatum* below.

### Multivoxel pattern of activity in dorsal anterior cingulate cortex predicts individual differences in context effect on probability estimate

We subsequently examined whether context effect on probability estimation can be represented in patterns of multivoxel activity. In our multivoxel decoding analysis, we tested whether we can predict individual differences in context effect based on patterns of multivoxel activity. A between-subject MVPA analysis was performed using a searchlight-based approach (Schmack et al., 2016). We analyzed multivoxel activity pattern separately at the time of stimulus presentation and reward feedback. We found that, at both time windows, multivoxel patterns of dACC activity significantly predicted subjects’ context effect at 50% reward (Fig. 7A-C), whereas multivoxel pattens of VMPFC activity predicted context only at the time of reward feedback (Fig. 7DE). For each subject separately, we computed the behavioral measure of context effect — the difference in mean probability estimate 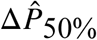 between the two different contexts: mean probability estimate of 50%-reward stimulus when it was in the [10%,50%] context minus the mean probability of the 50%-reward stimulus when it was in the [50%,90%] context). For each subject separately, we computed the neural measure of context effect — the difference in mean BOLD response to the 50% stimuli between the [10%,50%] context and [50%,90%] context, which we estimated for each voxel separately. In a searchlight (a sphere with 6mm radius) analysis, we trained a support vector regression (SVR) to learn and predict individual subject’s 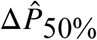 based on multivoxel pattern of neural measure within the searchlight and performed n-fold (n = 33 subjects) cross validation. The correlation between the left-out subjects’ 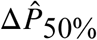 and the SVR-predicted 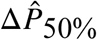 was then computed. To assess whether prediction performance was above chance, we used nonparametric permutation testing. For each searchlight, we constructed the null distribution of correlation coefficient by repeatedly doing the following: we randomly permuted the labels (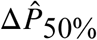) trained the SVR based on them, and computed the correlation between actual and predicted (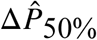). Prediction accuracy was considered significant if the probability of the true correlation was less than 0.05 Bonferroni-corrected for multiple comparisons across voxels in an *a priori* ROI. Here we considered three ROIs that have been shown to be involved in decision-making under uncertainty: medial prefrontal cortex (mPFC ROI based on Bartra et al., 2013), ventral striatum (VS ROI based on Clithero and Rangel, 2014), and dorsal anterior cingulate cortex (dACC ROI extracted from neurosynth.org). We found that both dACC and VMPFC showed significant prediction accuracy, but not VS. At stimulus presentation, 8 voxels in dACC survived Bonferroni correction for number of voxels tested in the ROI (magenta voxels in Fig. 7A); at reward feedback, 5 voxels at reward feedback survived Bonferroni correction (cyan voxels in Fig. 7A). The scatter plot in Fig. 7B shows the result from the peak voxel — searchlight centering on this voxel produces the largest correlation between predicted 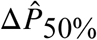 and actual 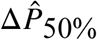 — at stimulus presentation (r=0.89, [x2,y30,z18]). The plots in Fig. 7C and 7E respectively show the results from the peak dACC voxel at reward feedback (r=0.83, [x2,y24,z20]) and peak VMPFC voxel at reward feedback (r=0.85, [x-12,y46,z0]). The dashed lines (Fig. 7BCE) represent perfect prediction, not regression fits.

**Figure 7.**
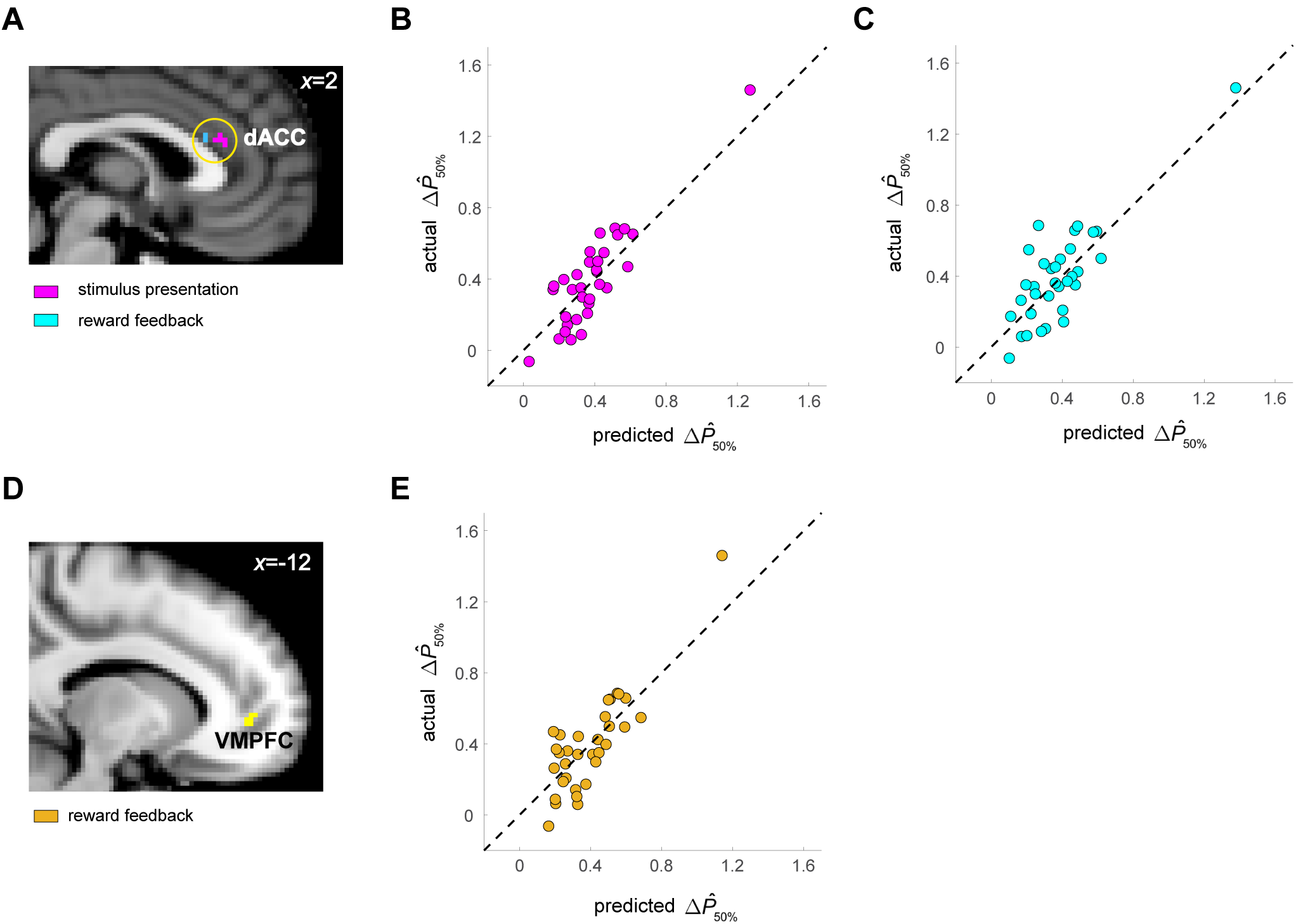
Dorsal anterior cingulate cortex (dACC) and ventromedial prefrontal cortex (VMPFC) represent context effect on probability estimate. **A-C.** dACC results. **A.** In a between-subject multivariate pattern analysis (MVPA), we found that patterns of multivoxel activity in dACC predicted context effect on individual subjects’ probability estimate (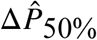) using activity patterns at the time of stimulus presentation (magenta) and at reward feedback (cyan) (p<0.05, Bonferroni corrected for 1350 voxels in the dACC ROI). **B.** We plot the actual 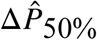 against predicted 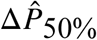 based on activity pattern — at the time of stimulus presentation — in the searchlight centered on the dACC voxel that produced the largest correlation between actual and predicted 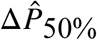 (referred to as the peak voxel). **C.** We plot the actual 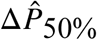 against predicted 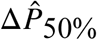 based on activity pattern — at the time of reward feedback — of peak voxel in dACC. **D-E.** VMPFC results. **D.** VMPFC voxels that significantly predicted behavioral context effect (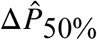) at the time of reward feedback (p<0.05, Bonferroni corrected for 1776 voxels in the VMPFC ROI). **E.** Scatter plot using data from the peak VMPFC voxel. Convention is the same as in **B-C.** The dashed line in **B,C** and **E** represents the 45-degree line, indicating perfect prediction.

### Superior temporal cortex and ventral striatum track reward statistics critical to context-dependent probability estimation

Based on univariate GLM analysis we found that, at stimulus presentation, the left superior to middle temporal cortex represented the frequency of reward associated with the stimulus (*f_S_*) and that the ventral striatum and fusiform gyrus represented the overall frequency of reward (*f*_overall_) (Fig. 8). These two statistics are critical to context-dependent probability estimation in our model. Our behavioral analysis indicated that *f_S_* was positively correlated with probability estimate, but *f*_overall_ was negatively correlated with probability estimate (see Fig. 6B). Our neural results also were consistent with these directions of correlation. We found that activity in the left posterior superior to middle temporal cortex (maximum z statistic = 3.91 at [x-64,y-36,z0] in the middle temporal gyrus) positively correlated with *f_S_* (Fig. 8A), while a distributed network including the ventral striatum (maximum z statistic = 4.31 at [x-18,y4,z-10]) and fusiform gyrus (maximum z statistic = 4.92 at [x34,y-70,z-18]) negatively correlated with *f*_overall_ (Fig. 8B; Table 1). By contrast, region-of-interest (ROI) analysis on VMPFC revealed no significant representation for both *f_S_* (t=0.56, df=33, p=0.579) and *f*_overall_ (t=-1.387, df=33, p=0.175). The VS ROI also did not significantly represent either *f_S_* (t=-1.259, df=33, p=0.217) or *f*_overall_ (t=-1.144, df=33, p=0.261) (Fig. 8C, beta plots on VMPFC and VS ROIs).

**Figure 8.**
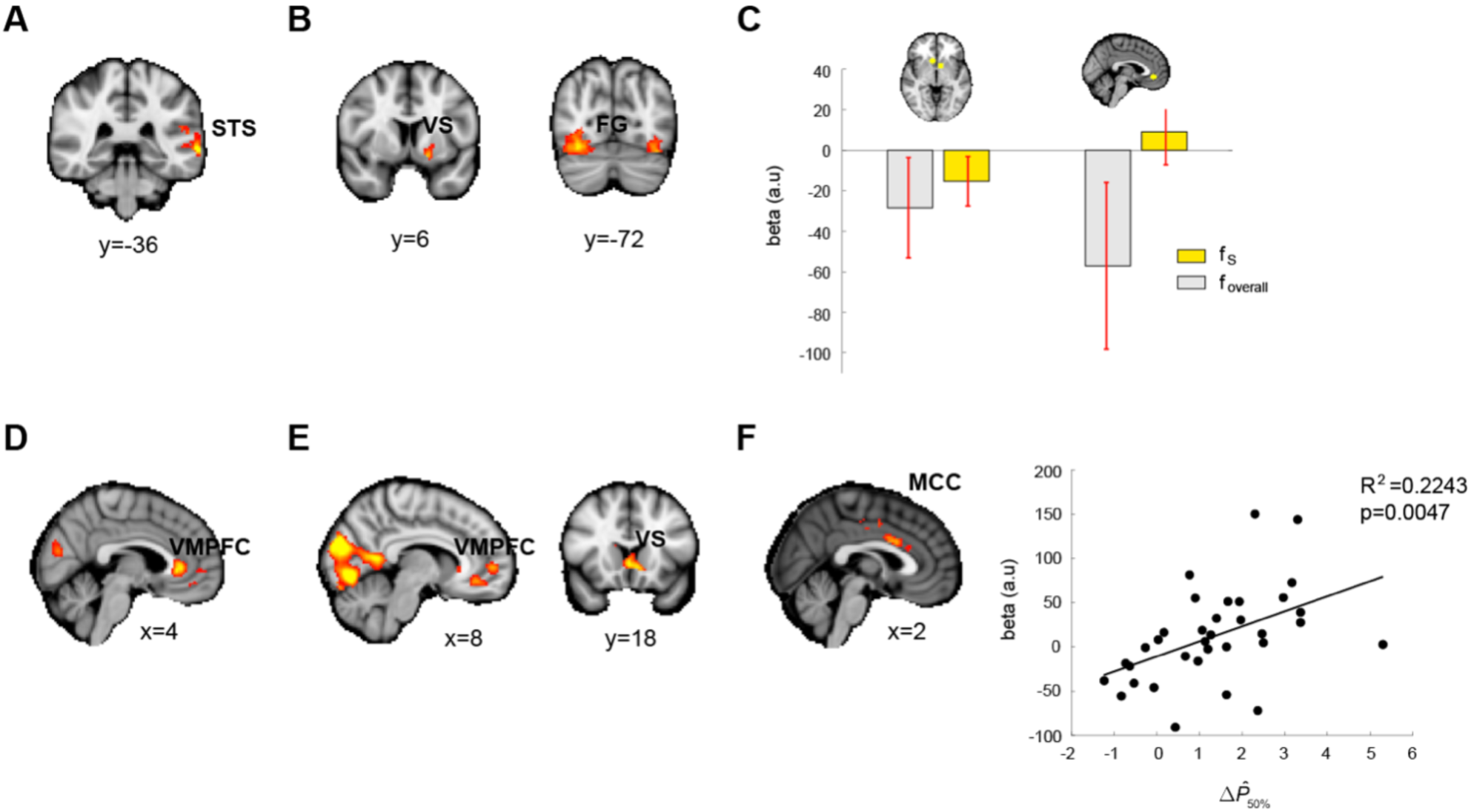
Univariate GLM results. **A-C.** Results obtained at the time of stimulus presentation. **A.** The left middle temporal gyrus (MTG) correlated with frequency of reward associated with stimulus (*f_S_*). **B.** The ventral striatum (VS) and fusiform gyrus (FG) correlated with the overall frequency of reward (*f*_overall_). **C.** Region-of-interest (ROI) analysis on medial prefrontal cortex (mPFC) and ventral striatum (VS) using coordinates from a meta-analysis paper (Clithero & Rangel, 2014). **D-F.** Results obtained at the time of reward feedback. Both regions did not significantly correlate with either *f_S_* or *f*_overall_. **D.** Reward-magnitude representation in VMPFC (whole-brain cluster corrected at p<0.05). **E.** Prediction-error signals in mPFC and VS (whole-brain cluster corrected at p<0.05). **F.** The middle cingulate cortex (MCC) represents individual differences in context effect on probability estimate (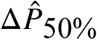) at the time of reward feedback in trials where subjects received a reward (whole-brain cluster corrected at p<0.05). The scatter plot is the result of leave-one-subject-out independent ROI analysis plotting estimated brain activity (beta, y-axis) against the behavioral measure of context effect (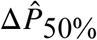) Each data point represents a single subject.

**Table 1:** Regions that significantly correlated with *f_S_* and *f*_overall_

| $f_S$ : positive correlation | | | | | |
| --- | --- | --- | --- | --- | --- |
|  | x | y | z | cluster size (voxels) | z statistic |
| Middle temporal gyrus (L) | -64 | -36 | 0 | 858 | 3.91 |
| $f_{\text{overall}}$: negative correlation | | | | | |

|  | x | y | z | cluster size (voxels) | z statistic |
| --- | --- | --- | --- | --- | --- |
| Fusiform gyrus (R) | 34 | -70 | -18 | 2556 | 4.92 |
| Fusiform gyrus (L) | -38 | -76 | -20 | 1429 | 4.43 |
| Putamen (L) | -18 | 4 | -10 | 1301 | 4.31 |

### Replication of reward prediction error signal and reward magnitude in VMPFC and striatum

Examining brain activity at the time of reward feedback, we replicated previous findings on reward magnitude (Bartra et al., 2013) and reward prediction error (McClure et al., 2003; O’Doherty et al., 2003; Abler et al., 2006; Rodriguez et al., 2006; Tobler et al., 2006; Hare et al., 2008) representations in the VMPFC and VS. Both VMPFC and VS represented reward magnitude (p<0.05 whole-brain FWE corrected for multiple comparisons using Gaussian random field theory with cluster-defining threshold at z>2.3) (Fig. 8D; Table 2) and reward prediction error, which is the difference between reward outcome (1=reward, 0=no reward) and subjects’ probability estimate (p<0.05 whole-brain FWE corrected for multiple comparisons using Gaussian random field theory with cluster-defining threshold at z>2.3) (Fig. 8E; Table 2). In a closer look at VMPFC representations for reward magnitude with higher threshold (z>3.6), we found two separate clusters – one in the medial OFC (maximum z statistic = 3.76, [x8,y44,z-12]) and one in the ACC (maximum z statistic = 4.31, [x2,y34,z2]). In addition, we also found that superior temporal gyrus, which we found to represent *f_S_*, also correlated with reward prediction error at feedback.

**Table 2.** Regions that significantly correlated with reward magnitude and reward prediction error at the time of feedback

| Reward magnitude: positive correlation |  |  |  |  |  |
| --- | --- | --- | --- | --- | --- |
|  | x | Y | z | cluster size (voxels) | z statistic |
| Visual cortex | -20 | -88 | -18 | 24282 | 6.33 |
| Anterior cingulate cortex | 2 | 34 | 2 | 2971 | 4.31 |

**Reward prediction error: positive correlation**
|  | x | Y | z | cluster size (voxels) | z statistic |
| --- | --- | --- | --- | --- | --- |
| Fusiform gyrus (R) | 36 | -68 | -18 | 16259 | 5.33 |
| Medial prefrontal cortex | -8 | 24 | -6 | 1936 | 3.89 |

At the time of reward feedback, we also found evidence for neural representations of context effect on probability estimate. In a group covariate analysis, for stimuli carrying 50% chance of reward that showed significant context effect, activity in the dACC extending to the mid-cingulate cortex (maximum z statistic = 3.32 at [x4,y14,z28]) correlated with individual differences in context effect (Fig. 8F; Table 3): the difference in activity between contexts in trials where subjects received a reward correlated with individual differences in 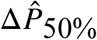 (p<0.05 whole-brain FWE corrected for multiple comparisons using Gaussian random field theory with cluster-defining threshold at z>2.3).

**Table 3:** Regions that significantly correlated with individual differences in context effect (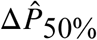)

|  | x | y | z | cluster size (voxels) | z statistic |
| --- | --- | --- | --- | --- | --- |
| Insula (L) | -60 | 2 | 8 | 912 | 3.46 |
| Midcingulate cortex | 4 | 14 | 28 | 683 | 3.32 |
| Insula (R) | 58 | 2 | 8 | 555 | 3.56 |

## DISCUSSION

This study was motivated by two observations on decision making under uncertainty. First, estimating probability of uncertain events is necessary to making good choices. In uncertain environments, events are probabilistic in nature and therefore having access to probability information is critical. In most cases, the decision maker is not given complete information about probability. Organisms must therefore estimate probability of events that are carriers of value in order to make good and adaptive choices. Second, context of experience should impact probability estimation. Since we rarely experience an event in isolation, the context of our experience – the presence of other events unfolding close in time – should play an important role in probability estimation. Here we found robust context effects on probability estimate: when a stimulus carried intermediate probabilities of reward (e.g. 50/50 of reward or no reward), subjects overestimated or underestimated its reward probability depending on whether the other stimulus present in the same context carried a smaller or larger probability of reward respectively. By contrast, subjects did not show significant context effect at extreme probabilities (e.g. 10% or 90% reward).

### Computational building blocks for context-dependent probability estimation

We developed a mathematical model for context-dependent probability estimation and tested whether it can describe the behavior observed in this study. In developing our model, we hypothesized that two computational building blocks – reference dependency and uncertainty dependency – play important roles in probability estimation. The first building block, reference dependence, describes that when estimating the probability of reward associated with a stimulus, the average probability of all stimuli a decision maker encounters in a context would serve as a reference point for comparison. Subjects would overestimate reward probability if the average probability were smaller than the stimulus reward probability. In contrast, if the average probability were larger than the stimulus probability, people would underestimate its probability of reward. Reference dependence has long been suggested in the evaluation of outcomes, treating outcomes as gains or losses relative to a reference point (Kahneman & Tversky, 1979) and reference point has been formally modeled as the decision maker’s expectation of future outcomes based on recent experience (Kőszegi & Rabin, 2006, 2007). However, there is very little work on examining whether probability estimation is reference-dependent (Otten & van der Pligt, 1996).

The second building block, uncertainty dependency, should only exist in estimation problems that are uncertain in nature. Given that there is more uncertainty regarding which potential outcome would occur when estimating probabilities that are near 0.5, they are more susceptible to context effect. This uncertainty dependency predicts that the reference-point effect on probability estimation would be modulated by uncertainty, i.e. variance or standard deviation of experienced outcomes associated with the stimulus of interest such that the effect is stronger at intermediate probabilities (larger standard deviation) than extreme probabilities (smaller standard deviation). These predictions are consistent with what we found in the current study: when stimulus reward probability was at the extremes (10% or 90% chance of reward), context did not significantly impact subjects’ probability estimate. By contrast, when a stimulus carried a 50% reward, subjects overestimated reward probability when the average reward probability in a context was smaller than 50% but underestimated reward probability when the average reward probability was larger than 50%.

The model we developed in this study and reinforcement learning (RL) models are not mutually exclusive. Subjects in our experiment had to learn probability of reward through feedback and it is highly likely that this learning process can be well described by RL models (Schultz et al., 1997; O’Doherty et al., 2003; Gläscher et al., 2010). What distinguishes our model from the standard model-free RL models is that, when estimating reward probability of a stimulus, our model explicitly states that not only is the learned frequency of reward associated with the stimulus is taken into account but also the estimated outcome uncertainty associated with it and the reward frequency of the other stimulus present in the same context. Having said that, it is possible to build and test a version of RL model that incorporates the context-dependent computations identified in this study and we anticipate that it would do reasonably well.

### Comparison between context-dependent probability estimation and subjective-value computations

This study was inspired by previous research that examined context effect on valuation but asked the question of whether and how context impacts probability estimation. While there are similarities between probability estimation and subjective-value computations, the results here highlight significant differences that are likely due to challenges imposed by estimating probability that are unique to valuation, which has been shown to be context-dependent (Tversky, 1969; Lichtenstein & Slovic, 1971, 1973; Grether & Plott, 1979; Slovic et al., 1988; Huber et al., 1982; Tversky & Simonson, 1993; Iyengar & Lepper, 2000; Kőszegi & Rabin, 2006, 2007; Abeler, J., Falk et al., 2011; Ericson & Fuster, 2011; Gill & Prowse, 2012). Recent neurobiological research also has begun to investigate how context impacts subjective-value representations in the brain (Tremblay & Schultz, 1999; Sallet et al., 2007; Elliot et al., 2008; Padoa-Schioppa, 2009; Kobayashi et al., 2010; Louie et al., 2011, 2013; Cox & Kable, 2014; Yamada et al., 2018) and how context-dependent value signals relate to choice behavior (Khaw et al., 2017; Zimmerman et al., 2018).

Here we discuss both similarities and differences between probability estimation and valuation, from the perspective of the tasks, computations, and algorithms carried out by the decision maker. First, while reward is present in both, a notable difference between valuation and probability estimation tasks is the presence or absence of uncertainty. The majority of valuation studies examined value computations in the absence of uncertainty (e.g. a stimulus is always associated with 3 drops of apple juice). In contrast, in our task subjects had to estimate the probability of receiving a reward associated with visual stimuli. Another difference in tasks is that while stimulus-reward associations are present in both, subjects’ own preferences for different rewards can clearly impact the subjective value of a stimulus but not its probability of reward. Instead, probability of an event such as reward is often determined by the outside world rather than subjective preferences that lie within an individual. Second, from the perspective of computations, by focusing on identifying computational building blocks for probability estimation and using them to guide model development, we were able to conclude that reference dependency is one crucial building block for both probability estimation and subjective-value computations. However, uncertainty dependency is unique to estimating rewards when there are uncertainties involved. Third, from the perspective of algorithms, our results suggest that probability estimation and subjective-value computations are carried out by different algorithms. This is supported by model comparison statistics showing that algorithms based on our context-dependent model (uncertainty-weighted reference-dependent model) performed better than normalization models that have been useful in describing subjective-value computations. In particular, uncertainty dependency is key to explaining both the observed context effect at intermediate probabilities and lack thereof at extreme probabilities. By contrast, normalization models – while it can explain context effect at intermediate probabilities – failed to describe non-significant context effect at extreme probabilities, especially at 90% reward. It is unclear how uncertainty statistics such as standard deviation of outcomes associated with a stimulus, which we found to be critical in explaining context effect on probability estimate, should be incorporated into the normalization models.

### Neural representations for context effect on probability estimation

Combining univariate and multivariate analyses, our fMRI results indicate that the neural implementations for probability estimation involve a network of brain regions, with dACC and VMPFC representing context effects on probability estimate and VS and superior temporal cortex representing reward statistics necessary for context-dependent probability estimation. The VMPFC finding, together with previous studies showing its involvement in context-dependent valuation (Tremblay & Schultz, 1999; Padoa-Schioppa, 2009; De Martino et al., 2009; Palminteri et al., 2015), established VMPFC as the common neurocomputational substrate for probability estimation and valuation. By contrast, the involvement of dACC may reflect aspects of probability estimation that is unique to valuation, which has been indicated in tracking uncertainty-related statistics such as variance and volatility (Behrens et al., 2007; Christopoulos et al., 2009), comparing recent and distant reward history (Wittman et al., 2016; Kolling et al., 2016), reinforcement-guided learning (Kennerley et al., 2006), and decision making under uncertainty (Rushworth & Behrens, 2008). Our results firmly established its involvement in context-dependent probability estimation, which requires computing reward-related statistics by tracking reward history.

If the dACC and VMPFC findings on context effect signified the output of context-dependent probability estimation, then the distinct regions in VS and superior temporal cortex we identified that tracked different reward statistics critical to probability estimation would highlight an initial stage for probability estimation. This pointed to the possibility that probability estimation is a two-stage computation that includes an initial stage of representing reward statistics followed by a second stage that utilizes these sources of information to compute probability estimate. Critically, we found that the ventral striatum, which connects with both dACC and VMPFC (Kunishio & Haber, 1994), represented the overall frequency of reward that serves as the reference point and that the superior temporal cortex, which had been shown to represent stimulus reward probability (Liljeholm et al., 2011; Tricomi & Lempert, 2015), represented stimulus reward frequency. Furthermore, there was behavioral-neural consistency in the direction of how these two reward statistics impacted probability estimate and neural activity: stimulus reward frequency had a positive correlation with both probability estimate and activity in the superior temporal cortex, while the overall reward frequency negatively correlated with both probability estimate and ventral striatum activity.

### Probability estimation and experience-based decision making

By showing that probability estimation is context-dependent, our results provide insights into experience-based decision making – an area of research that investigates decision making where choosers acquire information about different options through experience – in that context-dependent probability estimation is another source of bias that could affect people’s choice behavior. This research had shown that people nonlinearly weight probability information learned through experience (Hertwig & Erev, 2009) even when their probability estimates matched well with objective probabilities (Ungemach et al., 2009). However, context was typically not manipulated in these studies and therefore it is not possible to examine how context-induced bias on probability estimation would affect choice. It would be interesting to examine how the presence of these two sources of bias – probability weighting and context-dependent probability estimation – would impact choice behavior. At the neural algorithmic level, representations of probability distortion had been shown in the striatum (Hsu et al., 2009), lateral prefrontal cortex (Tobler et al., 2008), and VMPFC (Wu et al., 2011). Given that our results indicate that dACC, VMPFC, striatum and superior temporal cortex are critical to context effect, it remains open as to whether these two sources of biases are represented in the same brain regions and how they are being combined to impact experience-based decisions. In the future, it would also be important to examine, along with the stimulus standard deviation statistic, how other sources of variability in the environment could contribute to context effect. For example, how would the randomness or the rate of switching between different stimuli presented to the subjects influence context effect on probability estimate? How might this source of variability interact with the stimulus standard-deviation statistic to impact probability estimation?

In many choices we face, we are not explicitly given information about potential outcomes and their associated probabilities of occurrence. This makes probability estimation an essential computation for decision-making under uncertainty. Finding that probability estimation is context-dependent not only demonstrates that it is subjective, but more importantly that it is relative: subjective probability is not simply a representation of an abstract number. It reflects that seeing a stimulus with intermediate probability of reward like 0.5 feels better when the other stimulus present in the context is much worse than 0.5 than when the other stimulus carries a much better chance of reward. Such relative subjective probability is represented in the dorsal anterior cingulate cortex thought to be involved in extracting reward statistics from the environments in order to make decisions under uncertainty.

Probability estimation is a general problem organisms face in uncertain environments. Indeed, estimating probability through experience is an essential component of computational accounts of a wide array of cognitive tasks — from making sensory inferences, generating predictions to choosing between uncertain prospects (Tenenbaum et al., 2011; Pouget et al., 2013; Tonegawa et al., 2018). Previous work on statistical inference has developed and tested formal models for probability estimation but has often failed to consider how the impact of context, namely the presence of different events unfolding close in time, could impact probability estimation of each event of interest. Thus, the robust context effects we observed and the computational building blocks identified suggest that context may have strong effects on the many cognitive computations that require estimation of probability.

## MATERIALS AND METHODS

### Subjects

Thirty-seven right-handed subjects participated in the experiment (10 women, mean age = 22). Among them, 3 subjects did not complete the pre-fMRI session (see Procedure below). To be consistent in data analysis across subjects, these subjects’ behavioral and fMRI data were not analyzed. All participants gave written informed consent prior to participation in accordance with the procedures approved by the Taipei Veteran General Hospital IRB. All subjects received 500 NTD (1 US Dollar = 30 New Taiwan Dollar or NTD) for their participation and received additional bonus from the experiment. Their average earning was 1120 NTD.

### Task

We designed a simple stimulus-outcome association task to investigate how context affects probability estimation. Below we describe the trial sequence of the task and our manipulation of context. The task was programmed using the Psychophysics Toolbox in MATLAB (Brainard, 1997; Pelli, 1997). The stimuli were presented to the subjects through MRI-compatible goggles (Resonance Technology Inc.).

#### Trial sequence

On each trial, a red dot was first presented for 0.5s to indicate the start of a trial. This was followed by the presentation of a visual stimulus. Prior to the experiment, the subjects were instructed that each stimulus she or he faced carried a certain probability of reward. Upon seeing the stimulus, subjects had up to 2s to indicate his or her estimate of its reward probability with a button press. Failure to do so would prevent the lottery to be executed and as a result she or he won nothing on the current trial. There were 8 possible buttons each corresponding to a particular interval of probabilities: [0%,5%], [5%,20%], [20%-35%], [35%,50%], [50%,65%], [65%,80%], [80%,95%], [95%-100%]. Each button was assigned a finger to press. Subjects had to select the interval that best described his or her estimate of probability of reward associated with the current stimulus. Two opposite types of button assignments were implemented – left-to-right or right-to-right – and were balanced across subjects. In the left-to-right assignment, the probability increased from the left-most finger (left pinkie) to the right-most finger (right pinkie). The thumbs were not assigned. This design served to rule out confounds related to motor preparation and execution that would correlate with reward probability if not properly controlled. Once a button was pressed, subjects received visual feedback (0.5s) on the probability interval he or she indicated with the pressed button. A variable delay (1 to 7s) then followed before feedback on reward was provided (2s). The receipt or no receipt of a reward was determined by sampling with replacement from reward probability of the stimulus presented on the trial. If there was a reward, then the amount of reward was randomly selected (1 NTD to 5 NTD in steps of 1 NTD). In other words, reward magnitude was independent of reward probability. These design rules were explicitly stated to the subjects prior to the experiment. After the reward/no reward feedback, a variable inter-trial interval was presented (1 to 9s) before the next trial began.

#### Manipulation of temporal context

The subjects experienced three different contexts (context 1, 2, and 3) in the stimulus-outcome association task. Temporal context is implemented in a block of trials by placing two visual stimuli in it (Figure 1B). The order of their presentation was randomized for each block separately. Each stimulus in the context was assigned a unique probability of reward. On each trial, the subjects saw one of the two stimuli and had to sample from it in order to acquire information about its reward probability through feedback (see *Trial sequence* for details). Three reward probabilities, 10%, 50%, and 90%, were implemented in the task. Each probability appeared in two different contexts and was represented by two different visual stimuli. For instance, in both contexts 1 and 3, one of the two stimuli carries a 50% chance of reward. However, the reward probability of the other stimulus was different between these two contexts. In context 1, the other stimulus carried a 10% chance of reward. By contrast, in context 3, the other stimulus has a 90% chance of reward. This design therefore allowed us to compare two stimuli carrying the same probability of reward but were experienced in two different contexts where the other stimulus present carried different probabilities of reward. Specifically, we asked the questions – how would temporal context affect probability estimate on the stimuli carrying the same probability of reward? How might the effect of context change as a function of reward probability?

### Procedure

The experiment consisted of 3 distinct phases, pre-fMRI, fMRI, and post-test sessions. The experiment took approximately 60 minutes to complete.

#### Pre-fMRI session

Before the fMRI session, subjects first completed a behavioral session of the stimulus-outcome association task before entering the MRI scanner. There were three context blocks (1, 2, and 3 mentioned in *Task* above) of 40 trials each. In each block, there were 20 trials for each visual stimulus. The order of stimulus presentation in each block was randomized. This session served to train subjects to perform the task and to establish a certain degree of knowledge through experience about the reward probability of each stimulus under different contexts.

#### fMRI session

There were six blocks of trials in the session, two for each of the three temporal contexts. The ordering of the blocks was randomized for each subject separately. In each block, there were 30 trials, 15 trials for each of the two possible visual stimuli. The ordering of the stimulus presentation was randomized for each run separately.

#### Post-fMRI session

After the fMRI session, subjects performed a lottery choice task in which they were asked to choose between visual stimuli she or he faced in the previous two sessions. At this point, subjects should have acquired a certain degree of knowledge about the reward probability of each stimulus through experience. On each trial, the subjects were presented with two different visual stimuli. She or he was instructed that the amount of reward associated with winning was the same at 250 NTD across all stimuli and that she or he should choose the one preferred based on the experience with the stimuli, i.e. subjective belief about probability of reward. The subjects were told that at the end of the session one of his or her choices would be selected at random and realized. Since two visual stimuli were selected out of 6 stimuli on each trial, there were 15 possible pairs of visual stimuli. Among them, 3 pairs had stimuli with the same reward probability that were experienced under different contexts. For example, one pair had two visual stimuli both with 10% chance of reward. The 10% stimulus in context 1 was experienced along with a 50% stimulus. This 10% stimulus is referred to as the (10% | 10%-50%) stimulus. In contrast, the 10% stimulus in context 2 was experienced along with a 90% stimulus. This 10% stimulus is referred to as the (10% **|** [10%,90%]) stimulus. Besides the (10% | [10%,50%]) vs. (10% | [10%,90%]) pair, there were also the (50% | [50%,90%]) vs. (50% **|** [10%,50%]) pair, and the (90% | [10%,90%]) vs. (90% **|** [50%, 90%]) stimuli. The subjects’ choices in these 3 pairs thus allowed us to examine how context affected choice under each probability (10%, 50%, 90%) and compare context effect between these probabilities. In principle, there were 20 trials for each of these 3 pairs and 7 trials for the each of the remaining 12 pairs. Hence, there were a total of 144 trials. However, due to a programming error, not all pairs had the same number of trials: they ranged from 3 to 43 trials across subjects. The average number of trials for the 10%-reward, 50%-reward, and 90%-reward pairs was 27 (standard deviation = 7.64), 17 (standard deviation = 11.86), and 24 (standard deviation = 9.05) respectively (Fig. 3). We corrected this error in the behavioral control experiment (n=20 trials, for each pair on each subject; see *Behavioral control experiment* below) and replicated the choice result (Fig. 5D).

### Behavioral control experiment

One potential limit of the experiment is how we assigned buttons to the ranges of reward probability. The button that included 10% corresponded to the interval between 5% and 20%, the button that included 90% corresponded to the interval between 80% and 95%. It is possible that context also played a role in subjects’ probability estimates on the 10% and 90% stimuli, but as long as the difference in probability estimates between different contexts was not large enough such that estimates in the two contexts corresponded to two different buttons, we would not be able to detect it. In contrast, we had a more sensitive measure for context effect on the 50% probability stimuli because there were two buttons around 50%, one corresponding to the interval between 35% and 50% and the interval between 50% and 65%.

To rule out this potential confound, we conducted a behavioral control experiment by changing the button-probability assignment. In the control experiment (*n*=20 subjects), there were 10 keys, one corresponding to a 10% interval – [0,10%], [10%,20%], [20%,30%], [30%,40%], [40%, 50%], [50%,60%], [60%,70%], [70%,80%], [80%,90%], and [90%,100%]. For each stimulus, there were a total of 40 trials in the main session, which was equivalent to the fMRI session in the main experiment that had 30 trials for each stimulus. Otherwise, the control experiment was identical to main experiment.

### fMRI data acquisition

Imaging data were collected with a 3T MRI scanner (Siemens Magnetom Skyra) equipped with a 32-channel head array coil in the Taiwan Mind and Brain Imaging Center at National Cheng Chi University. T2*-weighted functional images were collected using an EPI sequence (TR=2s, TE=30ms, 35 oblique slices acquired in ascending interleaved order, 3.5×3.5×3.4 mm isotropic voxels, 64×64 matrix in a 224-mm field of view, flip angle 90°). Each subject completed 6 EPI blocks in the fMRI session. Each run consisted of 210 images. T1-weighted anatomical images were collected after the EPI runs using an MPRAGE sequence (TR=2.53 s, TE=3.3 ms, flip angle = 7°, 192 sagittal slices, 1×1×1 mm isotropic voxel, 256×256 matrix in a 256-mm field of view).

### fMRI preprocessing

The following pre-processing steps were applied using FMRIB’s Software Library (FSL) (Smith et al. 2004). First, motion correction was applied using MCFLIRT to remove the effect of head motion during each run. Second, we applied spatial smoothing using a Gaussian kernel of FWHM 5mm. Third, a high-pass temporal filtering was applied using Gaussian-weighted least square straight line fitting with *σ* = 50s. Fourth, registration was performed using a 2-step procedure. First, the unsmoothed EPI image that was the midpoint of the scan was used to estimate the transformation matrix (7-parameter affine transformations) from EPI images to the subject’s high-resolution T1-weighted structural image, with non-brain structures removed using FSL’s BET (Brain Extraction Tool). Second, we estimated the transformation matrix (12-parameter affine transformation) from the high-resolution T1-weighted structural image with non-brain structures removed to the standard MNI template brain.

### General linear modeling of BOLD response

**Model 1.** At the time of stimulus presentation, each visual stimulus (there were two in a black) was modeled by an indicator regressor whose length on each trial was the subject’s response time (RT) to indicate probability estimate. At the time of feedback, an indicator regressor, a regressor for the prediction error (PE), a regressor for the magnitude of reward outcome (MAG), and the interaction between PE and MAG were implemented.

#### Contrasts for examining contrast effect at stimulus presentation

We estimated the BOLD response to each stimulus and compared BOLD response between stimuli carrying the same reward probability but were experienced in different contexts. For each probability, at the subject level, we compared the BOLD response between two different contexts in a fixed-effect model. The resulting beta estimates were then fed into a group-level linear mixed-effect model to analyze whether there was a significant difference at the group level.

#### Contrasts for PE and MAG

After first-level time-series analysis for each block separately, beta estimates for PE and MAG were entered into a linear fixed-effect model for each subject. The results of the fixed-effect model on PE and MAG were separately entered into a group-level mixed-effect model to analyze the effect of PE and MAG at the group level.

#### Covariate analysis on context effect

This analysis focused on the 50%-reward stimuli that showed significant context effect on probability estimate. At the subject level, we separately estimated the BOLD response magnitude to the 50%-reward stimulus in the [10%,50%] 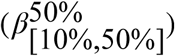context and in the [50%,90%] context (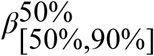). For each subject, we computed the difference between 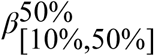 and 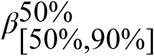, referred to as Δ*β*_50%_. We then performed a group-level covariate analysis on Δ*β*_50%_ (linear mixed-effect model) by using 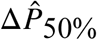 — the difference in the subject’s mean probability estimate (averaged across trials) on the 50%-reward stimulus — as a covariate.

**Model 2.** At stimulus presentation, we implemented three regressors, an indicator regressor and two parametric regressors. The length of these regressors is the subject’s trial-by-trial RT. One parametric regressor was the frequency of reward associated with the visual stimulus presented on the current trial (*f_S_*) and the other parametric regressor was the frequency of reward collapsed across different stimuli (*f*_overall_). Both were computed based on reward history in the past 10 trials. At the time of feedback, the regressors were identical to *Model 1*.

*Contrasts for f_S_ and f_overall_.* The analysis steps were identical to the one described *Contrasts for PE and MAG* above.

**Model 3.** At stimulus presentation, the regressors were identical to *Model 1*. At the time of feedback, we separately modeled each stimulus and reward outcome (reward or no reward).

#### Covariate analysis on context effect

This analysis was similar to the covariate analysis described in Model 1. We used 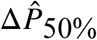 as covariate and separately analyze 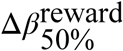 (for the trials in which subjects received a reward, the difference in BOLD response between different contexts for the 50%-reward stimuli) and 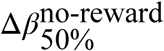 (for no-reward trials).

### Functional regions-of-interest (ROIs)

Three functional ROIs – the dorsal anterior cingulate cortex (dACC), ventromedial prefrontal cortex (VMPFC) and ventral striatum (VS) – were used. The dACC ROI was created using term-based meta-analyses (term: dorsal anterior) available in neurosynth.org. We used the reverse inference map generated from this analysis (299 studies) and excluded voxels in the map that are inconsistent with anatomical labeling of cingulate cortex. The resulting dACC ROI contained 1350 voxels (2mm isotropic). The VMPFC ROI (1838 voxels, 2mm isotropic) was created based on Bartra et al. (2013) that showed significant activation at the outcome receipt stage. The VS ROI (246 voxels, 2mm isotropic) was based on Clithero and Rangel (2014) that significantly correlated with subjective value.

### Between-subject multivoxel pattern analysis (MVPA)

The purpose of this analysis is to search for brain regions whose multivoxel pattern of activity can predict individual differences in context effect on probability estimate. Since context effect was significant when stimuli carried 50% chance of reward, the analysis focused on these stimuli. The behavioral measure of context effect is 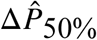 — the difference in probability estimate when 50%-reward stimulus was in the [10%,50%] context and in the [50%,90%] block — which we computed for each subject separately. At the neural level, for each subject separately, we obtained an Δ*β*_50%_ image as described in *covariate analysis on context effect above (Model 1 in General linear modeling of BOLD response)*. We adopted a searchlight procedure (Kriegeskorte et al., 2006) that allows the extraction of information from local pattern of multivoxel brain activity. We focused on the three ROIs — VMPFC, VS and dACC — described above (see *Functional regions-of-interest*). For a given voxel, we defined a spherical mask centered on it (radius = 8mm). We used 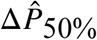 to label each of the multidimensional pattern vectors (vectors of multivoxel Δ*β*_50%_) and performed a leave-one-subject-out cross-validation to predict the normalized 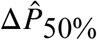. We excluded one subject from the analysis because she or he had 1/3 missing trials — trials in which no probability estimate was provided by the subject — for the 50% stimuli. Therefore, we performed the analysis on 33 subjects. We also performed the analysis by including all subjects and did not find the results to differ significantly (Fig. S1 in supplementary information). On each cross-validation run, we trained a linear support vector regression (SVR) machine with the labeled data from 32 subjects and tested on the independent data of the remaining subject. The SVR was performed using the LIBSVM implementation (http://www.csie.ntu.edu.tw/~cjlin/libsvm) with a linear kernel and a constant regularization parameter of c=1. Prior to SVR, both the continuous label (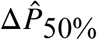) and the multidimensional fMRI pattern vector for a given voxel were normalized across subjects according to 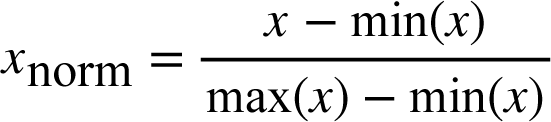 where *x* is the value before normalization, *x*_norm_ is the normalized value, max(*x*) min(*x*) max(*x*) is the maximum value of *x* and min(*x*) is the minimum value of *x*. On each cross-validation run, the normalization parameters (min(x), max(x)) were computed based on the training dataset and then applied to both the training and test dataset. This was to prevent the possibility that dependencies between test and training dataset were introduced. For each voxel, we obtained 33 predicted 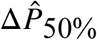 based on the above procedure. Prediction performance was determined by calculating Pearson’s correlation coefficient between the predicted and the actual 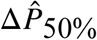. This procedure was repeated for all voxels.

To assess whether prediction performance was significantly above chance level, we used nonparametric permutation testing. For each voxel separately, we repeatedly trained and tested the SVR by randomly permuting the continuous label (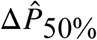) in order to generate a null distribution of the correlation coefficients. Prediction accuracy was considered significant if the probability of the true correlation occurring was P < 0.05, Bonferroni-corrected for multiple comparisons across all voxels in a priori regions-of-interest in the dACC (1350 voxels), VMPFC (1838 voxels), and VS (246 voxels). A whole-brain mask obtained from the univariate GLM analysis of this dataset was applied to these ROIs so that the voxels tested in these ROIs contained data from all subjects. As a result, the number of tested voxels in dACC, VMPFC and VS was 1350, 1776 and 202 respectively. The number of permutations run for each ROI was 1/(0.05/n) where n is the number of tested voxels. Therefore, the number of permutations run was 27000, 35520 and 4040 for the dACC, VMPFC and VS ROIs respectively. If the true correlation coefficient exceeded the largest value of the created null-distribution of correlation coefficients, the nonparametric P-value was reported as being smaller than the lowest possible nonparametric P-value given the number of permutations (1/*n*_permutations_). Note that even though the total number of possible permutations is fixed, this number is extremely large, which makes performing the analysis on all possible permutations very computationally expensive. This is why we used the approach described above, which is more realistic. In this case, the number of significant voxels (Fig. 7AC) could vary slightly each time one runs permutation test on the voxels within an ROI. However, the results reported in Fig. 7BCE survive even after we repeatedly performed permutation test.

### Modeling context-dependent computations for subjective probability: the uncertainty-weighted reference-dependent (VarRef) model

We developed a computational model – the uncertainty-weighted reference-dependent model (UncRef) – that takes into account temporal context and reward history. The subject’s probability estimate (*P_S_*) depends on two terms. First, it depends on the frequency of reward experienced in the recent past (*f_S_*). Second, it depends on a context term, which is the difference between *f_S_* and the overall frequency of reward (*f*_overall_) experienced in the recent past. Critically, the impact of the context term is determined by a parameter, which we refer to as the susceptibility parameter (). This is equivalent to saying that there is a gain control mechanism that regulates the impact of reference-dependent computation (*f_S_* − *f*_overall_) and that the susceptibility parameter represents the gain factor.

The model is expressed by the following equation

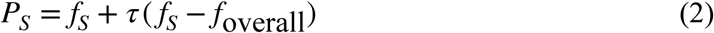

We assume that *τ* is sensitive to uncertainty on which potential outcome would occur, which can be defined by the variance 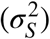 or standard deviation of experienced outcomes associated with the stimulus (*σ_S_*). Here, *τ* monotonically increases as a function of *σ_S_* and 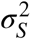.

In this experiment where subjects received binary reward outcomes (a reward or no reward), *σ_S_* should be the largest when the stimulus carried a 50% chance of reward. The model thus makes the prediction that the subjects’ probability estimate of the 50% stimulus is affected the most by (*f_S_* − *f*_overall_) than the other stimuli that carried either 10%-reward or 90% reward. The model also predicts that when a 50%-reward stimulus was experienced with a 10%-reward stimulus, subjects would overestimate reward probability, as (*f_S_* − *f*_overall_) would be greater than 0. In contrast, when the 50%-reward stimulus was experienced with a 90%-reward stimulus, subjects would underestimate reward probability (*f_S_* − *f*_overall_) < 0.

To directly test the model, we can implement *f_S_* and (*f_S_* − *f*_overall_) in a multiple regression model. Alternatively, we can also use *f_S_* and *f*_overall_ as the two regressors in the model. This model would have the slight advantage in that it reduced colinearity between the two regressors. Therefore, for each probability of reward (10%, 50%, 90%) separately, we performed the following regression analysis

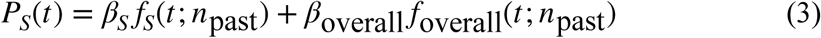

To make the computational model directly comparable to the regression model, we rewrite Eq. (2)

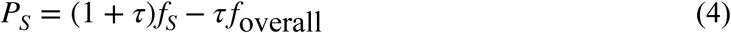

The model thus makes the following predictions on the regression coefficients: First, *β_S_* should be positively and significantly different from 0 across all reward probabilities. Second, *β_S_* associated with 50%-reward stimuli should be larger than that of the 10%-reward and 90%-reward stimuli. Third, *β*_overall_ associated with the 50%-reward stimuli should be negative and its magnitude should be larger than that of the 10%-reward and 90%-reward stimuli.

### Model fitting and model comparison

#### Computational models

A total of 11 different models – five versions of the UncRef model, three versions of the divisive normalization model (DivNorm) and three versions of the range normalization model (RangeNorm) – were fitted to the subjects’ probability estimates and were compared using model comparison statistic (Bayesian information criterion). The UncRef model takes the following form

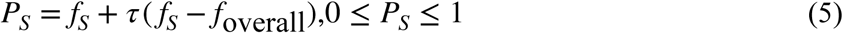

as described in Eq. (1) and we are explicitly modeling *τ* by the standard deviation of reward outcome in the recent past (*τ* = *σ_S_*) (UncRef-sigma) or variance of past reward outcomes 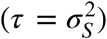 (UncRef-var). Here we used the past 10 trials to calculate *f_s_*, *f*_overall_, 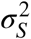, *σ_S_*.

To further model overestimation of small probabilities and underestimation of large probabilities, which we observed in our dataset and was also found in Fox & Tversky (1998), we use the following equation to model the these estimation biases

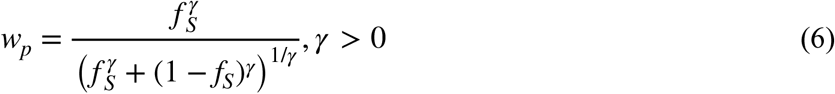

where *w_p_* represents the biased probability estimate that overestimate small probabilities and underestimate large probabilities when *γ* <. Then the probability estimate is computed by the same UncRef principles

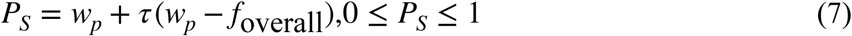

The models that replace *f_S_* with *w_p_* are labeled as the UncRef-sigma-wp and UncRef-var-wp models. We also fitted a version of the model by replacing *f*_overall_ with *f*_OS_ (frequency of reward associated with the other stimulus in the same context) and used *σ_S_* as a measure of *τ* (UncOS-sigma).

The divisive normalization model (DivNorm) takes the following form

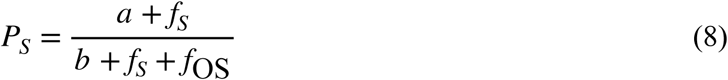

where *f*_OS_ is the other stimulus in the same context as *f_S_*. For example, in a [10%,50%] context, if *f_S_* corresponds to the stimulus carrying 50% chance of reward, then *f*_OS_ would be the stimulus carrying 10% reward. This form is referred to as the DivNorm-3params as there were three parameters in the model – two from Eq. (8) and one noise parameter described in Eq. (10). We also fitted a two-parameter version of the divisive normalization model (Louie et al., 2011) (DivNorm-2params: parameter *b* in the denominator in Eq. (8) and the noise parameter *σ*_noise_ in Eq. 10) and a one-parameter version of the model (DivNorm-1param: taking out parameters *a* and *b* in Eq. (8) and having only *σ*_noise_ in Eq. 10).

The range normalization model (RangeNorm) takes the following form

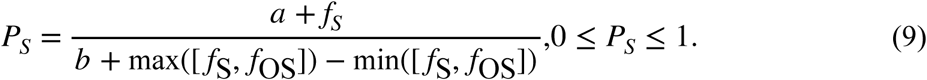

This form is referred to as the RangeNorm-3params as there were three parameters in the model –two from Eq. (9) and the noise parameter *σ*_noise_ in Eq. (10). We also fitted a two-parameter version of the range normalization model (RangeNorm-2params: parameter *b* in the denominator in Eq. (9) and the noise parameter *σ*_noise_ in Eq. 10) and a one-parameter version of the model (RangeNorm-1param: taking out parameters *a* and *b* and having only *σ*_noise_ in Eq. 10). In Table 4, we summarize the 11 different models and their respective free parameters. Note that the parameter *σ*_noise_ (Eq. 10) is not considered a free parameter of these models. Rather, it is free parameter for modeling the stochasticity of probability estimate *P_S_* once it is being computed in these models.

**Table 4:**
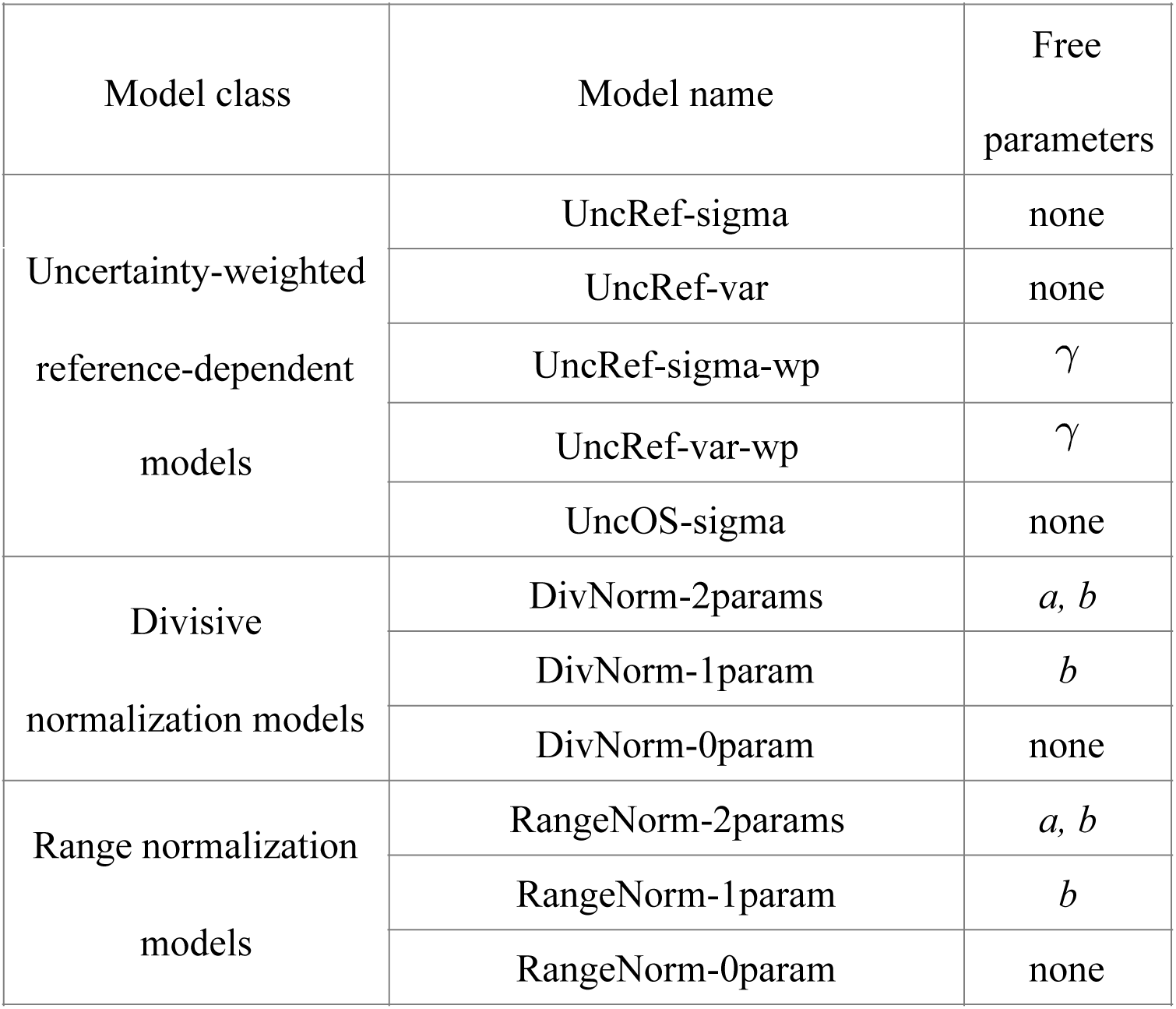
Models and their free parameters

#### Maximum-likelihood estimation

We computed the mean probability estimates (averaged across all subjects) time-series data (the one shown in Figure 5A-C) and fitted it with the four computational models described above. We assume that probability estimate is a Gaussian random variable with mean at 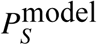 and variance 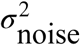 where 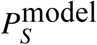 is the model-predicted probability estimate and 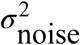 is a free parameter. The likelihood function is therefore

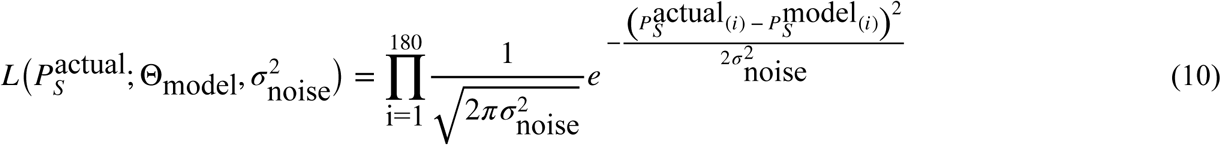

where 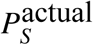 is a vector containing the subjects’ mean probability estimates, Θ_model_ represents the set of free parameters in a model, and 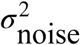 is the variance of the Gaussian noise (a free parameter).

For example, for the UncRef-sigma model, Θ_model_ is empty because there is no free parameter in it and therefore there is only one free parameter (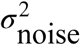) to be estimated when fitting UncRef-sigma.

#### Model comparison

For each model separately, we used non-parametric Bootstrap method to reconstruct the distribution of Bayesian information criterion (BIC) described below

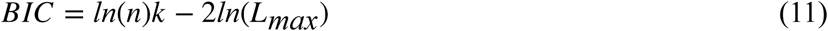

where *n* represents the number of trials (*n*=180), k represents the number of free parameters in a model and *L*_max_ represents the value of maximum likelihood. We resampled the subjects with replacement 10,000 times to construct 10,000 resampled dataset. For each resampled dataset, we computed the mean probability estimate (across subjects) associated with each trial in the order. We then fitted the probability estimates with the model and based on the fitting result computed BIC. As a result, for each model separately, we obtained a distribution of BIC values, which we used to compute 95% confidence interval and to compare between different models.

## ACKNOWLEDGMENTS

This work was supported by the Ministry of Science and Technology in Taiwan (MOST 104-2410-H-010-002-MY3 and MOST 106-2420-H-010-003) and Brain Research Center, National Yang-Ming University from The Featured Areas Research Center Program within the framework of the Higher Education Sprout Project by the Ministry of Education (MOE) in Taiwan to SWW.

## SUPPLEMENTARY INFORMATION

MVPA analysis using all subjects’ data. Detailed analysis steps are described in the main text. In the main text (Fig. 7), we present results that excluded one subject’s data because she or he had too many missing trials (1/3 of the total trials in the 50% condition where she or he did not provide probability estimate within two seconds after stimulus onset), making estimates of BOLD response less reliable compared with other subjects. In the results shown below (Fig. S1), we included this subject’s data in the analysis. As expected, the results are not identical to those shown in Fig. 7. However, they are similar in the sense that dACC — at both stimulus presentation and reward feedback — represents individual subjects’ context effect on probability estimate, while mPFC at reward feedback represents context effect.

**Figure S1.**
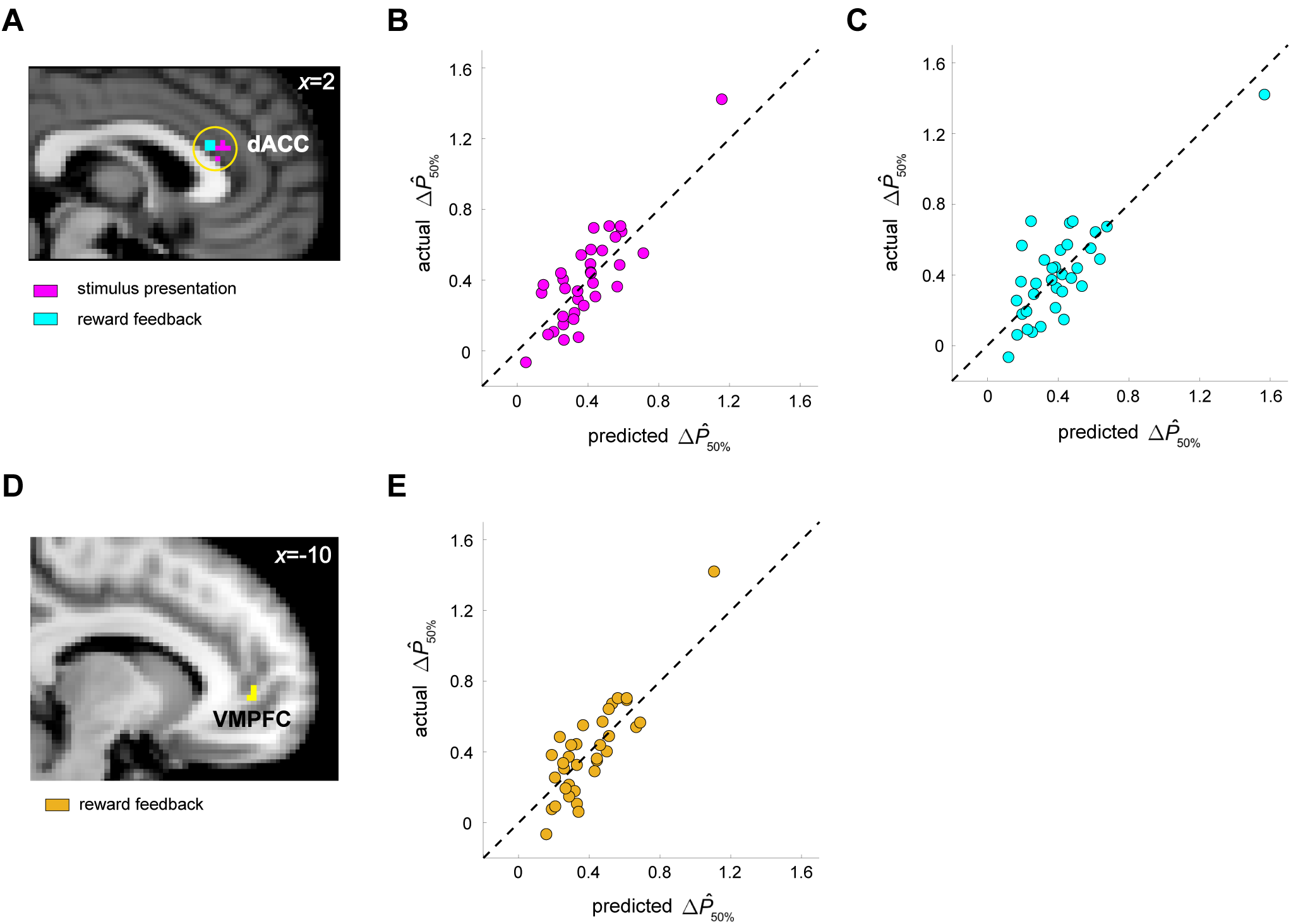
Dorsal anterior cingulate (dACC) and ventromedial prefrontal (VMPFC) activity predict context effect on probability estimate. **A-C.** dACC results. **A.** In a between-subject multivariate pattern analysis (MVPA), we found that patterns of multivoxel activity in dACC predicted context effect on individual subjects’ probability estimate (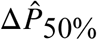) using activity patterns at the time of stimulus presentation (magenta) and at reward feedback (cyan) (p<0.05, Bonferroni corrected for 1350 voxels in the dACC ROI). **B.** We plot the actual 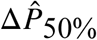 against predicted 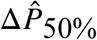 based on activity pattern — at the time of stimulus presentation of the peak voxel in dACC. The dotted line represents 45-degree line, indicating perfect prediction. **C.** We plot the actual 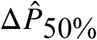 against predicted 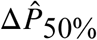 based on activity pattern — at the time of reward feedback of the peak voxel in dACC. **D-E.** VMPFC results. **D.** VMPFC voxels (1776 voxels in VMPFC ROI) that significantly predicted behavioral context effect (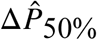) at the time of reward feedback. **E.** Scatter plot using data from the peak VMPFC voxel. Convention is the same as in **B-C**. The dashed line in **B,C** and **E** represents the 45-degree line, indicating perfect prediction.

